# TCF4 mutations disrupt synaptic function through dysregulation of RIMBP2 in patient-derived cortical neurons

**DOI:** 10.1101/2023.01.19.524788

**Authors:** Brittany A. Davis, Huei-Ying Chen, Zengyou Ye, Isaac Ostlund, Madhavi Tippani, Debamitra Das, Srinidhi Rao Sripathy, Yanhong Wang, Jacqueline M. Martin, Gina Shim, Neel M. Panchwagh, Rebecca L. Moses, Federica Farinelli, Joseph F. Bohlen, Meijie Li, Bryan W. Luikart, Andrew E. Jaffe, Brady J. Maher

## Abstract

Genetic variation in the transcription factor 4 (*TCF4)* gene is associated with risk for a variety of developmental and psychiatric conditions, which includes a syndromic form of ASD called Pitt Hopkins Syndrome (PTHS). *TCF4* encodes an activity-dependent transcription factor that is highly expressed during cortical development and in animal models is shown to regulate various aspects of neuronal development and function. However, our understanding of how disease-causing mutations in TCF4 confer pathophysiology in a human context is lacking. Here we show that cortical neurons derived from patients with *TCF4* mutations have deficits in spontaneous synaptic transmission, network excitability and homeostatic plasticity. Transcriptomic analysis indicates these phenotypes result from altered expression of genes involved in presynaptic neurotransmission and identifies the presynaptic binding protein, RIMBP2 as the most differentially expressed gene in PTHS neurons. Remarkably, TCF4-dependent deficits in spontaneous synaptic transmission and network excitability were rescued by increasing RIMBP2 expression in presynaptic neurons. Together, these results identify TCF4 as a critical transcriptional regulator of human synaptic development and plasticity and specifically identifies dysregulation of presynaptic function as an early pathophysiology in PTHS.

## Introduction

Transcription factor 4 (TCF4) is a basic helix-loop-helix (bHLH) transcription factor that is genetically associated with a variety of neuropsychiatric disorders including schizophrenia, bipolar disorder, post-traumatic stress syndrome, major depressive disorder, and autism spectrum disorder (ASD) ^1–5^. Rare, autosomal dominant mutations in transcription factor 4 (TCF4) are causal for Pitt Hopkins syndrome (PTHS), a syndromic form of ASD often associated with intellectual disability, developmental delay, absent or limited speech, motor delay, constipation, facial features and in some cases breathing abnormalities and/or seizure activity ^6–10^ Given the clinically pleiotropic nature of TCF4, gaining insight into the neurobiology of TCF4 and how its dysregulation can lead to pathophysiology is critical to improving our understanding of disease mechanisms related to psychiatric disorders and will eventually inform the development of therapies.

TCF4 is highly expressed during cortical development where it forms dimers with itself or other bHLH transcription factors (e.g. NeuroD1, Id2, Ascl1, Olig1) and regulates target gene expression ^11^. The peak of TCF4 expression coincides with neurogenesis and neuronal migration, but continues to be expressed at lower levels throughout gliogenesis and into adulthood ^12,13^. Cell-type specific expression analysis showed that TCF4 is expressed across all the major cell types of the cortex, including excitatory and inhibitory neurons, astrocytes and oligodendrocytes ^14.^ Consistent with this expression pattern, TCF4 is shown to regulate many aspects of cortical development, excitatory neuronal function, and oligodendrocyte development ^12,13,15–19^. In addition, TCF4 is known to regulate transcription in response to neuronal activity, and was shown to play a role in long-term potentiation ^15,20–22.^

Despite evidence of conservation in *TCF4* expression across species ^12,13,16^, very little is known about *TCF4* function in the human context. PTHS causing mutations can result in either TCF4 haploinsufficiency or can lead to the expression of dominant-negative and/or hypomorphic protein which contain either a nonfunctional bHLH domain or activation domain ^23–25^. To improve our understanding of the role of TCF4 in neuronal development and function, we modeled human cortical development in PTHS syndrome using patient-derived induced pluripotent stem cells (hiPSCs) with mutations that result in nonfunctional bHLH domains. All of our PTHS patient lines showed consistent physiological deficits in their membrane properties, intrinsic excitability, spontaneous synaptic transmission and network activity. In addition, PTHS neurons had decreased magnitude of homeostatic plasticity and dysregulation in both activity-dependent and developmental transcription related to synaptic transmission. Importantly, we identified RIMBP2 as a critical presynaptic protein that is dysregulated in PTHS patient lines and we demonstrate that viral expression of RIMBP2 in presynaptic neurons effectively rescued spontaneous synaptic transmission and network activity deficits.

## Results

### Characterization of PTHS and control 2D cortical cultures

hiPSCs were generated from four PTHS patient lines harboring missense mutations in the bHLH domain and two patient lines harboring nonsense mutations, along with six neurotypical controls (Figure 1A). hiPSCs were differentiated into forebrain specific neurons using dual SMAD inhibition by adapting previously published methods (Figure 1B) ^26,27^. At DIV25, NPCs were dissociated and plated onto rat primary astrocytes and allowed to differentiate, leading to cortical co-cultures that were previously shown to have improved neuronal maturation ^26,28^. Cortical fate-specification at 6 weeks (wks) was assessed by staining for canonical markers of lower layer (TBR1, CTIP2) and upper layer (SATB2, CUX1) excitatory neurons (Figure 1C). We calculated the percentage of each layer marker based on the total number of human nuclei (hNA+), and observed that mutations in TCF4 had no effect on the percentage of TBR1+, CTIP2+, CUX+, or SATB2^+^ cells (Figure 1D). As previously described, approximately 10% of the neurons were GABAergic (GAD65/67+) ^26,27^, with similar proportions of inhibitor neurons found in both control and PTHS cultures (Supp Figure 2A,B).

**Figure 1.**
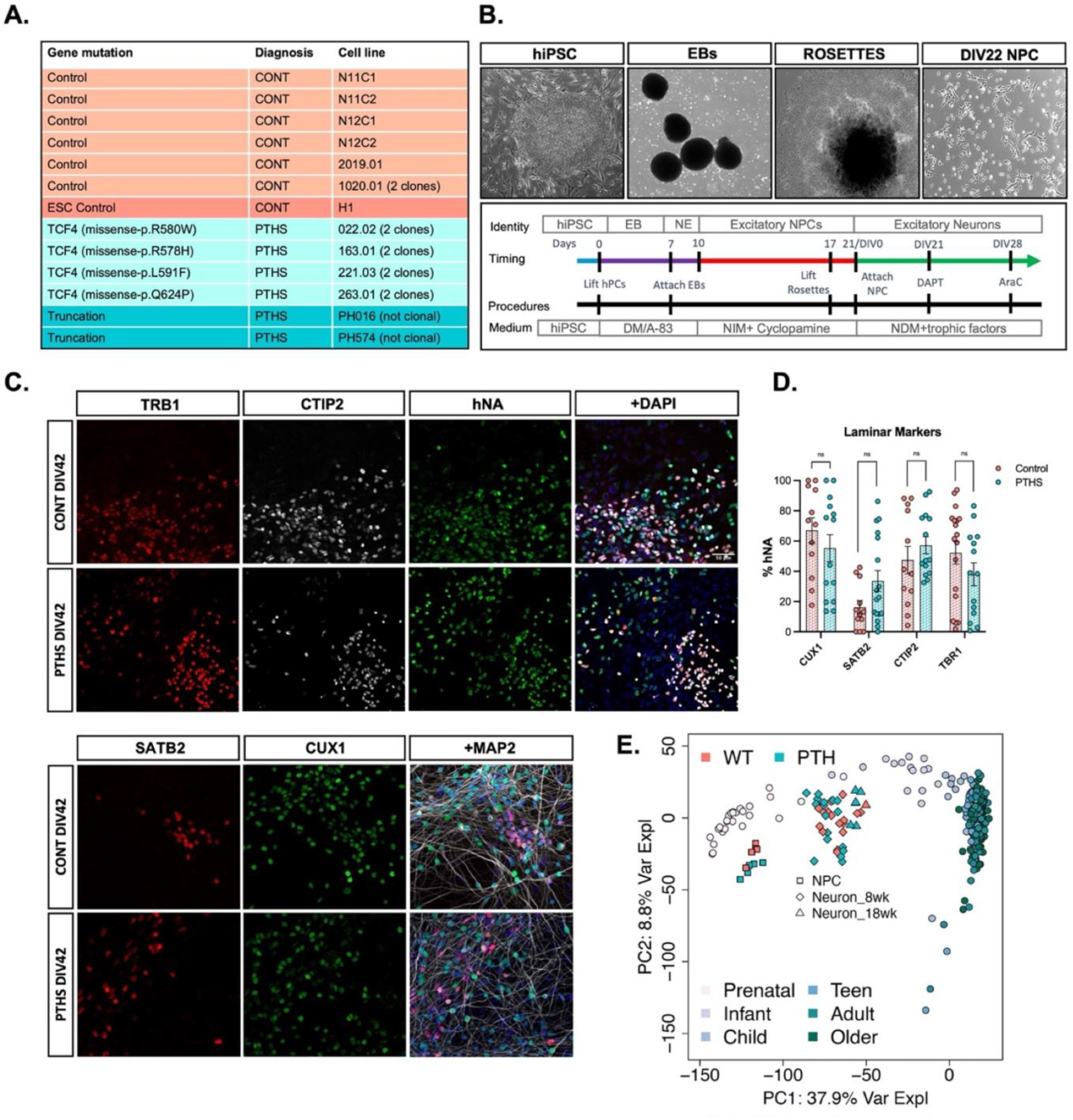
2D differentiation of PTHS patient-derived hiPSCs generated cortical neurons of both upper- and lower-layer fates. **A**. Human control and PTHS cell line information. **B**. Diagram of the cortical differentiation protocol showing example live cell images of hiPSC, embryoid bodies (EBs), neural rosettes, and neural progenitor cells (NPCs) and schematic of cell identity, timing, procedures, and cell culture media. **C**. Representative images of cortical neurons immunostained for various cortical layer markers. Upper panel, ICC staining for lower layer markers CTIP2 (L5, some L2-4) and TBR1 (L5-6). Lower panel, ICC staining for upper layer markers SATB2 (L2-4, some L5) and CUX1 (L2-3, some human NPCs). **D**. Quantification of layer specific markers showing diagnosis has no effect on fate specification. Each data point represents the percentage of hNA co-labelled with a layer-specific marker, averaged across three fields per experiment, for each line; from four independent differentiations. **E**. PCA plot comparing the developmental time course of our cortical differentiation protocol to development of the human DLPFC.

**Figure 2.**
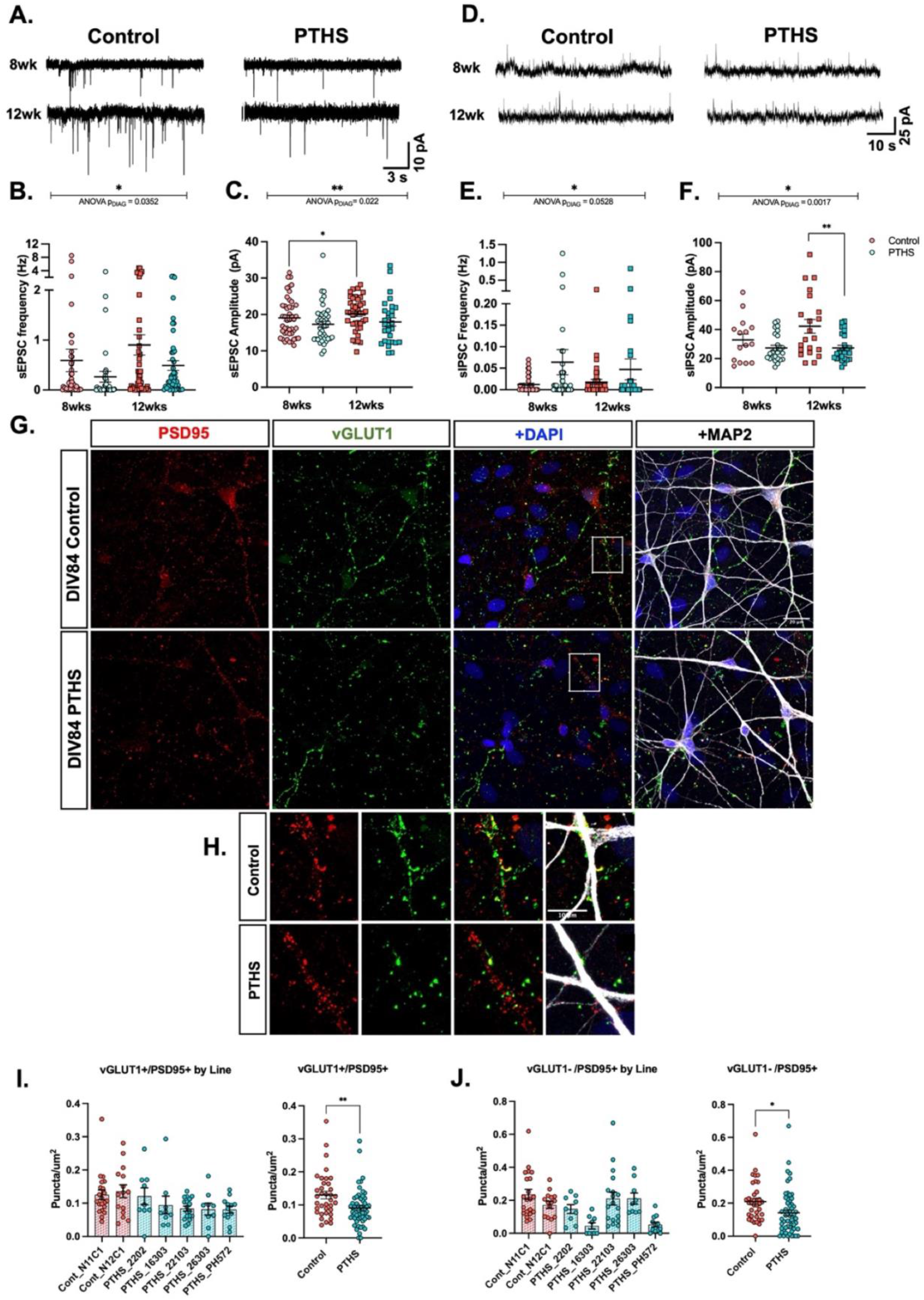
PTHS patient-derived neurons displayed altered spontaneous synaptic transmission and reduced glutamatergic synaptic density. **A**. Example traces of sEPSCs recorded at -70mV from 8wk and 12wk control and PTHS neurons. **B**. Summary data showing a main effect of diagnosis on the sEPSC frequency F (1, 175) = 4.503, p = 0.0352. **C**. Summary data showing a main effect of diagnosis on the sEPSC amplitude (F (1, 153) = 5286, p = 0.022). **D**. Example traces of sIPSCs recorded at 0mV from 8wk and 12wk control and PTHS neurons. **E**. Summary data showing a marginal effect of diagnosis on sIPSC frequency (F (1, 159) = 3.806, p = 0.0528). **F**. Summary data showing a main effect of diagnosis on sIPSC amplitude (F (1, 82) = 10.58, p = 0.0017), which was driven by reduced sIPSC amplitudes in PTHS neurons at 12wks (*t* (1,182) = 3.504, p = 0.0015). Each data point in **B, C, E, F**. represents individual neurons from five control lines and four PTHS lines, from a minimum of four independent differentiations, collapsed by diagnosis, see Supp Figure 7 for individual line data. **G**. Representative ICC images showing staining for excitatory synapses containing PSD95+/VGLUT1+puncta on MAP2+ dendrite in 12wk control and PTHS cortical neurons (20µM scale). **H**. Inlay from white box in **G**. showing zoomed in images of excitatory synaptic puncta (10µM scale). **I**. Summary data showing PTHS neurons had significantly fewer PSD95+/vGLUT1+ puncta compared to controls (*t* (1,91) = 3.080, p = 0.0027), a pattern consistent across all PTHS lines. **J**. Summary data showing PTHS neurons had significantly fewer PSD95+ only puncta compared to controls (*t* (1, 91) = 2.506, p = 0.014), a pattern consistent across all PTHS lines. Each data point in **I-J**. represents the average of three independent neurite segments, which in sum are >300µm in length, averaged per field; three fields per line, per experiment, across a minimum of three independent differentiations.

To further validate cell identity and to quantify developmental progression of our cortical cultures we performed transcriptional analysis on RNA collected from NPCs (DIV0) and neurons at 8 wks and 18 wks. We compared gene expression between our cortical cultures and human postmortem dorsolateral prefrontal cortex (DLPFC) samples across the lifespan (Figure 1E). We observed the largest source of variation (PC1: 37.9%) corresponded to cell state/development. Our 8wk cultures approximated a late prenatal stage of development, while 18wk cultures begin to overlap early stages of infancy (Figure 1E). In addition, cellular deconvolution was used to analyze the fraction of RNA expressed by specific cell types. As expected, there was a significant developmental decrease in the fraction of NPC RNA in 8wk and 18wk cultures, a developmental increase in the fraction of neuronal RNA, and there was no difference in these measures due to diagnosis (Supp Figure 1B-C). Similarly, no differences were associated with the fraction of RNA associated with either glial or endothelial sources by diagnosis (Supp Figure 1D-H). These results indicate that our cortical differentiation protocol recapitulates gene transcription observed in the developing human DLPFC with our neuronal co-cultures approximating the third trimester and into early infancy.

Next, we performed differential gene expression analysis by diagnosis, developmental time and the interaction between diagnosis and time (Supp Table 1). We identified 40 DEGs (Benjamini-Hochberg corrected p-value, “P_adj”<0.05) by diagnosis and 12718 DEGs (P_adj<0.05) due to developmental time. Gene set enrichment analysis of the DEGs by diagnosis (p<0.005) identified dysregulation of several gene ontology (GO) terms broadly related to development and indicative of the expression of TCF4 in a variety of tissues ^29–31.^. Some example GO terms included embryonic development, cell fate commitment, and the development of striated muscle tissue, renal system, reproductive system, heart, epidermis and the ear (P_adj<0.05; Supp Table 2). These pathways were all upregulated in PTHS cells, suggesting that in the neural lineage TCF4 represses expression of non-neural lineage specific genes. Together, these results indicate that mutations in TCF4 disrupts expression of genes related to development, however this altered expression does not appear to disrupt the overall cellular composition of the cortical cultures derived from PTHS patient hiPSCs compared to controls.

### TCF4 mutations delay the maturation of membrane properties and intrinsic excitability in cortical neurons

To analyze the effect of *TCF4* mutation on physiological development, we performed whole-cell electrophysiological recordings in 8 wk and 12 wk neurons and measured developmental changes in the neuronal membrane properties and intrinsic excitability. Over this 4 week period, all cortical cultures showed stereotypical maturation, as we observed significant decreases in membrane resistance, hyperpolarization of the resting membrane potential, increases in Na^+^ current density, and increases in the maximum acceleration of action potentials (APs; Supp Figure 3-6). However, effects of diagnosis were observed, as PTHS neurons displayed a significant increase in membrane resistance and a reduction in K^+^ current density compared to control neurons (Supp Figure 3D, I). These diagnosis effects were driven by a greater difference at the 8 wks compared to 12 wks, and suggests a developmental delay in aspects of intrinsic excitability and membrane properties in PTHS neurons.

**Figure 3.**
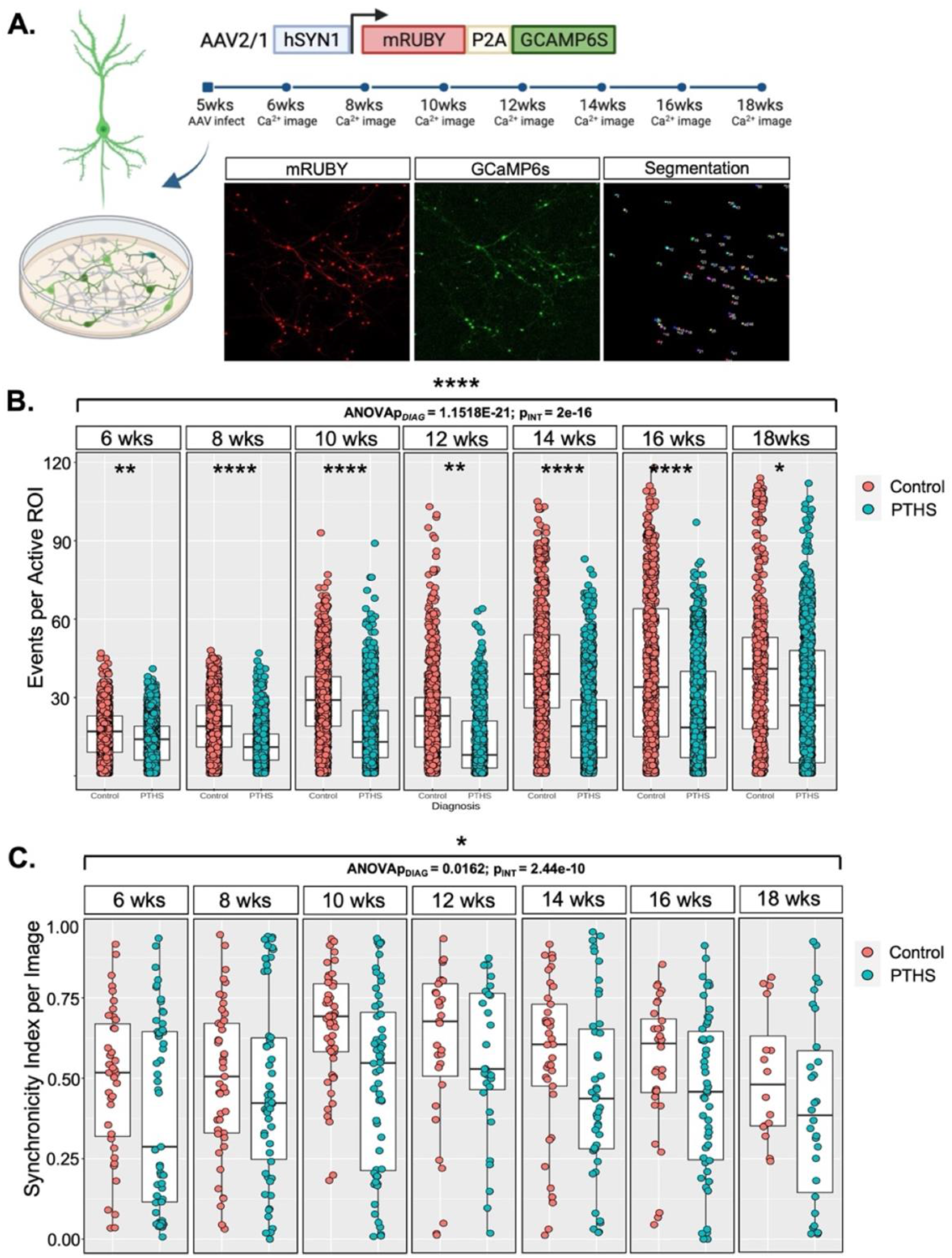
PTHS patient-derived neurons displayed decreased spontaneous network activity throughout development. **A**. Schematic showing the experimental paradigm and representative images of viral infected neurons co-expressing mRuby and GCaMP6s. The mRuby signal was used for segmentation and generation of regions of interest (ROIs) used for quantification of fluorescence changes due to Ca2+ influx. **B**. Summary plot showing a main effect of diagnosis on the number Ca^2+^events across active ROIs between 6wks and 18wks (diagnosis effect = -9.318, p = 1.2e-21) and a main interaction between diagnosis and age (INT effect = 15.541, p < 2e-16). Post hoc analysis shows a significant diagnosis effect was observed at each time point (6wks, diagnosis effect = -0.2079, p = 0.0119; 8wks, diagnosis effect = -0.3889, p = 2.22e-05; 10wks, diagnosis effect = -0.5382, p = 1.02e-08; 12wks, diagnosis effect = -0.5785, p = 0.0021; 14wks, diagnosis effect = -0.4814, p = 3.74e-05; 16wks diagnosis effect = -0.5636, p = 0.0001; 18wks, diagnosis effect = -0.5010, p = 0.0354). Post hoc analysis of the interaction between diagnosis and age show effects at a subset of time points (10wks, INT effect = -7.885, p = 0.0142; 14wks, INT effect = -13.427, p < 0.00014; 16wks, INT effect = -13.076, p = 0.0002; 18wks, INT effect = -8.576, p = 0.043), which is not shown in the figure. Each data point represents an active ROI from an individual neuron. Developmental assay videos were recorded from seven control lines and ten PTHS lines, four point mutants with two clones each and two truncated lines are represented; each line was recorded from two to four differentiations every other week. **C**. Summary plot showing a main effect of diagnosis on the synchronicity of Ca^2+^ events between 6wks and 18wks (diagnosis effect = -0.1018, p = 0.0162). A significant interaction effect between diagnosis and age was also observed (INT effect = 0.3959, p = 2.44e-10). Each data point represents an individual field, two fields were recorded per line, per week, from each differentiation run.

There were no within line differences across any of these measurements in the control neurons (Supp Figure 3-6). However, there was variability within the PTHS lines, including differences in membrane capacitance (Supp Figure 4E) and within line differences in Na^+^ and K^+^ current densities (Supp Figure 5B, E), AP acceleration (Supp Figure 6B, E), and maximum AP output (Supp Figure 4B). Post hoc analysis identified a single line in each of these measurements that was responsible for the within diagnosis differences. Importantly, when we removed the line driving the within diagnosis effect for each measure, all main effects of diagnosis remained significant, thus indicating the diagnosis effect was not being driven by any one PTHS line (Supp Figure 4C, 5C, 5F, 6C, 6F).

**Figure 4.**
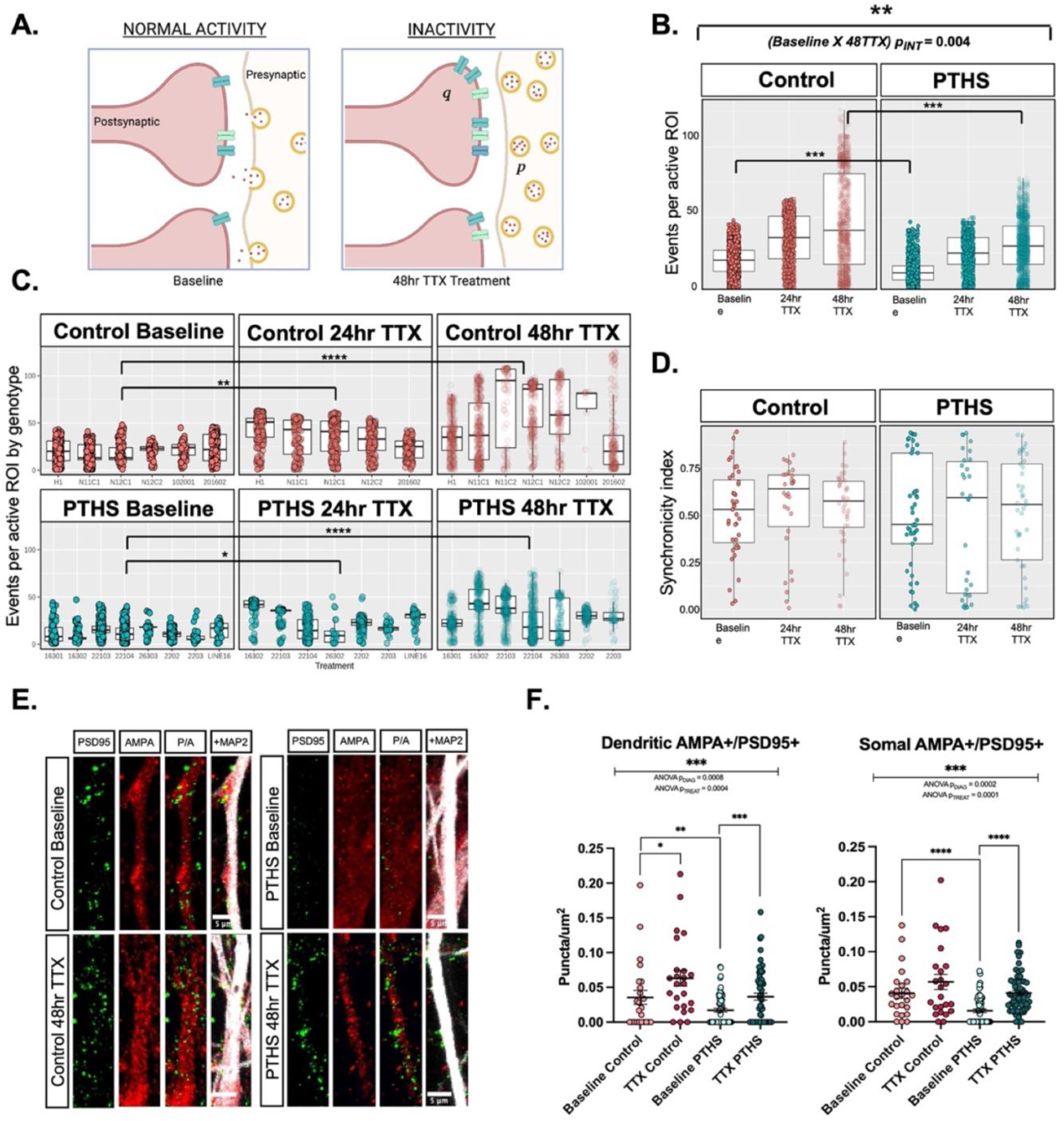
PTHS patient-derived neurons displayed abnormal homeostatic plasticity. **A**Schematic of synaptic upscaling after prolonged inactivity. After blocking synaptic transmission via 48hrs of TTX, upscaling can occur after postsynaptic changes in quantal amplitude via multiplicative changes in glutamate receptor number (***q***) and/or presynaptic changes in vesicle size, number or release probability (***p***). **B**. Summary data showing the magnitude of network upscaling (events per active ROI) after 48hr TTX treatment is significantly reduced in PTHS neurons compared to controls (PTHS effect = -11.94, p = 0.0046). Post hoc analysis confirmed PTHS neurons showed a reduced number of events per active ROI compared to control neurons (p = 0.0006) and following 48hr TTX treatment (p = 0.0005). Each data point represents individual ROIs from 6-7 control lines and seven PTHS lines, with one field recorded for each condition, across >2 differentiations per line, with a total of 4 differentiations. **C**. Summary data by line showing significant network upscaling (events per active ROI) across control lines after 24hr (treatment effect = 0.3977, p = 0.0022) and 48hr (treatment effect = 0.8515, p = 2.75e-18) TTX treatment. A similar effect is observed in PTHS patient lines (24hr TTX, treatment effect = 0.5838, *p* = 1.78e-09; 48hr TTX, treatment effect = 0.2852, p = 73.38e-03). Each data point represents individual ROIs recorded above in B., but represented by line. **D**. Summary data showing TTX treatment had no effect on the network synchronicity index. Each data point represents an individual field, one field was recorded per line, per condition, per week, from a minimum of two differentiations per line, with a total of four differentiations. **E**. Example ICC image showing staining of AMPA receptors, PSD95 and MAP2 expression in control and PTHS neurons at baseline (top) and after 48hr TTX treatment (bottom). **F**. Summary data showing a significant effect of treatment on the density of AMPA+/PSD95+ puncta on dendrites (F (1, 164) = 12.83, p = 0.0004) and soma (F (1, 164) = 15.49, p = 0.0001). A significant effect of diagnosis on the density of AMPA+/PSD95+ puncta on dendrites (F (1, 164) = 11.80, p = 0.0008) and soma (F (1, 164) = 14.84, p = 0.0002). Each data point in **E**. and **F**. represents the average of three independent neurite segments, which in sum are >300µm in length, averaged per field; four fields per line, per experiment, across three independent differentiations.

**Figure 5.**
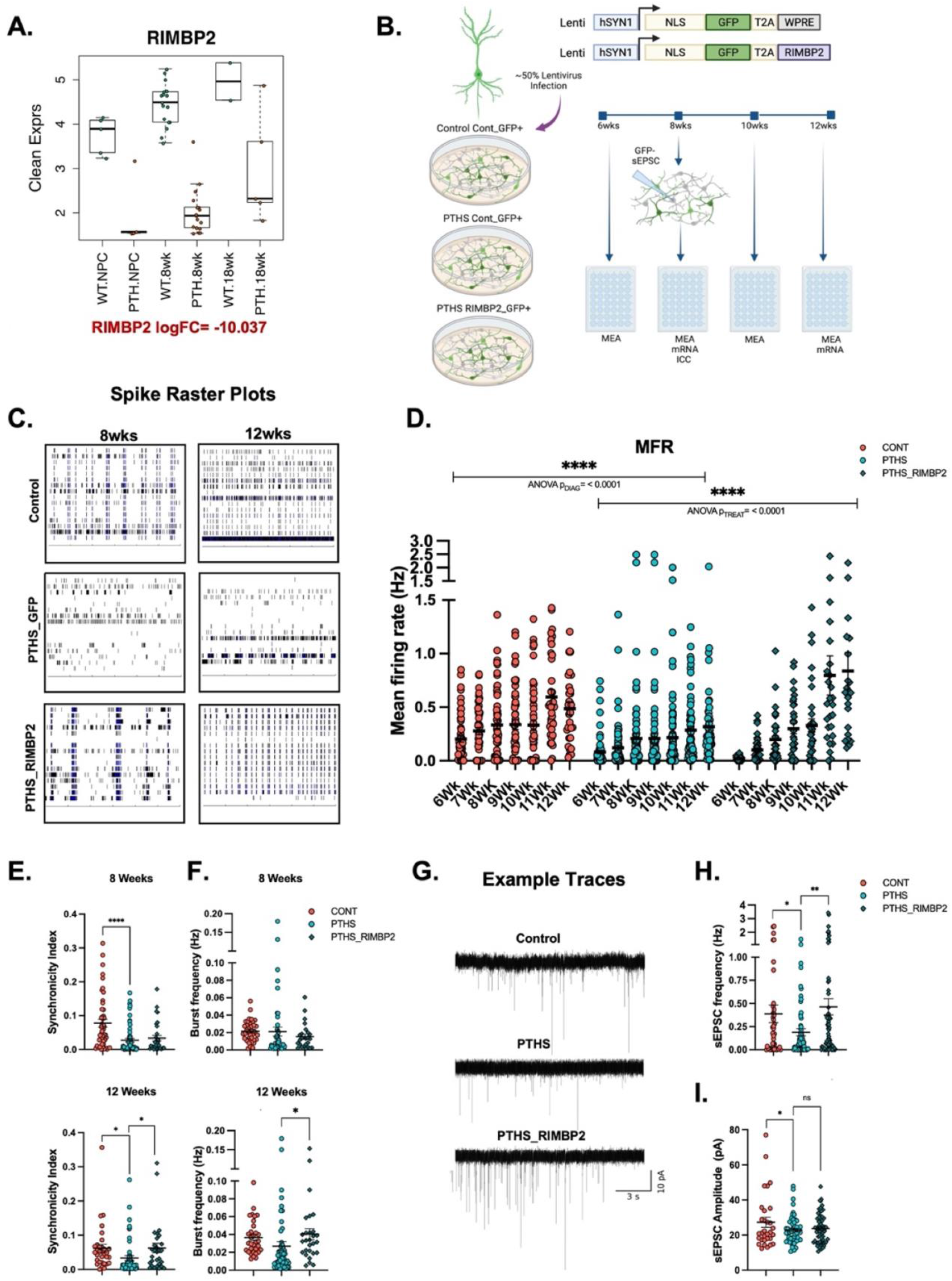
Presynaptic expression of RIMBP2 rescued spontaneous network activity in PTHS patient-derived cortical neurons. **A**. Summary plot showing a main effect of diagnosis on RIMBP2 expression (logFC = -10.04, P_adj = 1.72e-4). **B**. Schematic describing RIMBP2 rescue experiments: ∼50% of the cultures were infected at 4wks with lentivirus constructs; MEA recordings were carried out weekly from 6-12wks; at 8wks patch clamp electrophysiology was carried out on non-transduced postsynaptic cells and both mRNA and ICC were conducted. **C**. Representative raster plots demonstrating MEA activity at 8 and 12wks of recording. **D**. Summary graph showing a main effect of diagnosis on the mean firing rate (MFR, F (1, 810) = 47.95, p < 0.0001) and a main effect of age (F (6, 810) = 11.21, p < 0.0001). A significant main effect of viral treatment on MFR in PTHS patient neurons (F (1, 706) = 29.55, p < 0.0001) and a significant interaction between treatment and time (F (6, 706) = 8.640, p < 0.0001). **E**. Summary plot showing a diagnosis effect on the MEA synchronicity index at 8wks (*t* (1, 108) = 4.339, p < 0.0001) and 12wks (*t* (1, 84) = 2.229, p = 0.0142). RIMBP2 expression in PTHS neurons increased synchronicity by 12wks (*t* (1, 81) = 2.125, p = 0.0183). **F**. Summary plot showing RIMBP2 expression in PTHS neurons increased MEA burst frequency by 12wks (*t* (1, 76) = 1.637, p = 0.0529). Each data point in **D**-**F**. represents a single well, from wells with greater than two active electrodes, from two control lines and five PTHS lines, across three independent differentiations, with two or more wells per condition, per differentiation. **G**. Representative traces of sEPSCs recorded from control, PTHS, and RIMBP2 treated (PTHS_RIMBP2) neurons at 8wks. **H**. Summary plot showing a diagnosis effect on the baseline sEPSC frequency at 8wks (*t* (1,105) = 2.197, p = 0.0150) and a significant effect of RIMBP2 treatment (*t* (1,135) = 2.728, p *=* 0.0036). **I**. Summary plot showing a diagnosis effect on the baseline sEPSC amplitude at 8wks (*t* (1, 82) =1.719, p = 0.0447). Each data point in **H**. and **I**. represents individual neurons from two control lines and four PTHS lines collapsed by diagnosis from a minimum of three differentiations.

A developmental delay in PTHS neurons was also reflected in the ability of PTHS neurons to generate trains of APs, a phenotype that was previously reported in rodent models ^16,22^Control neurons reliably generated trains of APs in response to current steps (Supp Figure 3A, B). However, PTHS patient neurons showed a significant reduction in the number of APs generated in response to current pulses (Supp Figure 3A, B) and a decrease in the maximum AP output (Supp Figure 3F) at 8wks, which was resolved by 12wks. These results suggest that disease-causing mutations in TCF4 lead to delays in the maturation of neuronal properties essential to the development of their intrinsic excitability, but by 12wks, most measurements of membrane properties and intrinsic excitability are not different by diagnosis.

### Reduced spontaneous network activity in PTHS neurons

We next assayed spontaneous synaptic transmission in our neuronal cultures. Spontaneous excitatory (sEPSCs) and inhibitory postsynaptic currents (sIPSCs) were recorded in individual neurons by holding the cell at -70mV and 0mV, respectively. PTHS neurons had significant reductions in sEPSC frequency and amplitude compared to controls (Figure 2A-C), an effect that was consistently observed across all lines (Supp Figure 7A-D). Control neurons also demonstrated a developmental increase in sEPSC amplitude between 8wks and 12wks that was not observed in PTHS neurons (Figure 2C). PTHS neurons had a modest increase in sIPSC frequency, but reduced sIPSC amplitude, which was predominantly driven by the 12wk time point (Figure 2D-F). Again, this effect was consistently observed across all lines (Supp Figure 7E-H). These results suggest that the balance between excitation and inhibition is altered during network development in PTHS patient neurons compared to control.

To investigate cellular mechanisms associated with these spontaneous transmission deficits, we quantified the density of excitatory and inhibitory synaptic puncta in 12wk cultures. We observed a significant decrease in the density of both excitatory (VGLUT1+/PSD95+; Figure 2G-K) and inhibitory synaptic puncta (GAD65&67+/Gephyrin+; Supp Figure 2A,C,D) in PTHS neurons compared to controls. Thus, the observed decrease in the sEPSC frequency and sIPSC amplitude in PTHS neurons is in part explained by an overall reduction in synaptic connectivity, and suggests that TCF4 regulates synaptic development and connectivity in human neurons.

### Reduction in network-wide Ca^2+^ transients in PTHS neurons throughout development

Next, we investigated whether the observed deficits in synaptic transmission and connectivity at the cellular level disrupted function at the network level. Network activity was measured using a genetically encoded Ca^2+^ indicator, GCaMP6s, to chronically monitor changes in Ca^2+^ influx from 6wks to 18wks of development. In a prior study, we demonstrated this spontaneous network activity in human neurons was AP-dependent, as acute treatment with tetrodotoxin (TTX) completely abolished network activity ^26^. Consistent with the diagnosis effects on sEPSC frequency, PTHS neurons showed a significant reduction in the frequency of AP-dependent spontaneous Ca^2+^ transients and a significant interaction between diagnosis and time, indicating network development was slower in PTHS neurons (Figure 3B). We also observed a significant reduction in the proportion of active neurons in a field (Supp Figure 8B) with no diagnosis effects on the number of neurons per field (Supp Figure 8A). Lastly, we quantified the global synchronicity index for each image (Supp Figure 8C, D) and observed PTHS cultures showed a significant reduction in network synchronicity (Figure 3C).

Given that both inhibitory and excitatory transmission were disrupted in PTHS neurons (Figure 2), we further investigated the role of these neurotransmitters on network activity. Acute treatment of cultures with the GABAa receptor antagonist picrotoxin (PTX; 100µM) had no effect on the frequency of Ca^2+^ events in either control or PTHS cultures (Supp Figure 8E). However, when cultures were acutely treated with the glutamate receptor antagonists, DNQX and AP5, the vast majority of network activity was abolished, independent of diagnosis (Supp Figure 8F). All together, these results suggest that disease-causing mutations in *TCF4* significantly reduce the development of spontaneous network-wide activity, which appears to occur primarily through disruptions in spontaneous excitatory synaptic transmission.

### Decreased magnitude of synaptic upscaling and reduction in global AMPA puncta

Prior studies indicate that TCF4 is an activity-dependent transcription factor that regulates long term potentiation ^15,20–22^; however, TCF4 regulation of homeostatic plasticity has not been established. Chronic treatment with TTX is known to robustly trigger synaptic homeostasis through upscaling of both pre- and postsynaptic function and transcription (Figure 4A) ^32–35^. Therefore, we examined homeostatic plasticity and downstream activity-dependent transcription in response to chronic inhibition of network activity in PTHS and control cultures. We treated GCaMP6s infected cultures at 8wks with TTX for 24 and 48 hours (hrs). In control neurons, both 24hr and 48hr TTX treatment resulted in a significant increase in the frequency of Ca^2+^ events compared to baseline control cultures (Figure 4B), demonstrating for the first time, that homeostatic upscaling is robustly measurable in human neurons, via network-wide Ca^2+^ imaging. We also observed network upscaling in PTHS patient lines, however PTHS neurons displayed a significant reduction in the magnitude of upscaling compared to controls (Figure 4B), a finding that was consistent between PTHS lines (Figure 4C). TTX treatment showed a trend for increased synchronicity, however no differences were associated with diagnosis (Figure 4D), and there were no differences in the number or proportion of active neurons within a field after treatment (Supp Figure 9A, B).

As presynaptic and postsynaptic scaling effects are indiscernible in Ca^2+^ imaging, we next addressed whether differences in the magnitude of upscaling were associated with the accumulation of postsynaptic AMPA-type glutamate receptors, as previously reported ^35,36^. At baseline, control neurons demonstrated significantly greater numbers of AMPA+/PSD95+puncta around the soma and along dendrites compared to PTHS neurons (Figure 4E, F), which further supported the observed reductions in spontaneous synaptic transmission (Figure 2A-C), synapse density (Figure 2G-I), and network activity (Figure 3A, B). Both control and PTHS patient neurons demonstrated significant treatment effects in response to 48 hr TTX treatment, as we observed increased AMPA+/PSD95+ puncta around the soma and along dendrites (Figure 4F). However, PTHS neurons showed a reduction in the magnitude of upregulation of AMPA+/PSD95+ puncta along the soma and dendrites compared to control neurons (Figure 4E, F). This diagnosis effect was consistent with the reduced magnitude of network upscaling (Figure 4B). Together, these data indicate that PTHS neurons display synaptic upscaling, but *TCF4* mutations appear to limit the magnitude of this homeostatic response.

To evaluate transcription in response to synaptic upscaling, we collected RNA from baseline and 48hr TTX-treated cultures from both PTHS and control neurons and performed RNA sequencing. In control cultures, TTX treatment resulted in differential expression of 49 genes (P_adj<0.05; Supp Table 3). Gene set enrichment analysis of DEGs (p<0.005) in control cultures identified a variety of pathways associated with the glutamatergic synapse, synaptic vesicles, presynapse, ion channels, learning and memory, and synaptic plasticity (Supp Figure 10A, Supp Table 4). When comparing these 49 DEGs to previously published DEGs in response to synaptic upscaling in primary rat cultures ^37^, we observed approximately 40% overlap in DEGs between human and rat neurons (Supp Figure 10B, Supp Table 5). Additionally, a number of genes previously identified as essential for synaptic plasticity and homeostasis were represented (e.g. *ACTL6B, CPLX1, CAMK2B, GABRA5, KCNF1, KCNJ3, KCNN1, MAPK8IP1, NCS1, NGEF, SYNGR1, SYP, VGF*), suggesting human and rodent neurons elicit a similar transcriptional response to homeostatic upscaling ^37–42^.

In stark contrast to control neurons, TTX treatment of PTHS neurons did not result in any DEGs after FDR correction (P_adj<0.05; Supp Table 6), an effect that may explain the observed diagnosis effect on the magnitude of upscaling (Figure 4B, F). These results highlight the importance of activity-dependent transcriptional regulation by TCF4 to appropriately regulate synaptic and network homeostasis.

### Increased RIMBP2 expression drives recovery of intrinsic excitability and network-wide activity in PTHS neurons

To investigate potential mechanism(s) responsible for the reduction in spontaneous network activity in PTHS patient neurons, we next analyzed transcriptional changes due to diagnosis by combining RNA sequencing data obtained from 8wk and 18wk neurons. In total, 162 DEGs survived FDR correction (P_adj<0.05) (Supp Table 7). Gene set enrichment analysis of these DEGs (p<0.005) identified downregulation of a surprising number of early developmental and cell division associated pathways (Supp Table 8). Importantly, we also observed downregulation in pathways associated with the presynaptic terminals and regulation of neurotransmission (Supp Figure 11A; Supp Table 8), which corresponds well with observed phenotypes. Numerous DEGs in these pathways are essential regulators of presynaptic vesicle machinery (e.g. *CACNB2, STX1A, STY1, UNC13C* and *RIMBP2*), which could underlie deficits in the frequency of spontaneous synaptic transmission and network activity. In fact, the top DEG in PTHS cultures across development was Rab-interacting molecule-binding protein 2 (*RIMBP2*), a presynaptic protein that regulates the probability of synaptic transmitter release at central synapses (Figure 5A, Supp Table 1) ^43–45^. Interestingly, two prior ChIP-seq studies identified TCF4 binding to the *RIMBP2* gene locus in rat NPCs and in human SH-SY5Y cells ^16,46^. These ChIP-seq studies in combination with the observation that *RIMBP2* expression is consistently downregulated across developmental stages (Figure 5A; NPC and neurons) suggests TCF4 directly regulates RIMPB2 expression.

The dysregulation of RIMBP2 expression and its role in presynaptic transmission led us to reason that reduced RIMPB2 in PTHS patient neurons could underlie the observed deficits in synaptic transmission and network excitability. To test this molecular mechanism, we infected PTHS patient neurons with a lentivirus that expresses either RIMBP2 plus GFP or GFP alone under the control of the hSYN1 promoter at 5wks (Figure 5B) ^44^. We observed that lentiviral infection resulted in GFP expression in approximately 50% of the neurons, which was independent of the lentiviral construct or cell line (Supp Figure 12A, B) To confirm that viral transduction resulted in the up-regulation of *Rimbp2* expression, we performed qPCR in 8wk and 12wk cultures, using a mouse specific probe for *Rimbp2*. We detected a marginal increase in lentivirus driven mouse *Rimbp2* expression at 8wks, which reached significance by the 12wks in cultures with 50% transduction efficiency (Supp Figure 12C).

Next, we tested the effectiveness of *Rimbp2* expression at rescuing network-wide activity deficits in PTHS neurons using multi-electrode arrays (MEAs). Neurons were grown on MEAs and infected with lentivirus at 5wks and spontaneous network activity was monitored between 6wks and 12wks (Figure 5B). The GFP alone lentivirus had no effect on the mean firing rate (MFR) compared to uninfected cultures regardless of diagnosis (Supp Figure 13A, B), and therefore uninfected and GFP alone infected control wells were collapsed by line in our subsequent analysis. For both PTHS and control lines, MFR significantly increased over development which was independent of diagnosis (Supp Figure 13C). Consistent with Ca^2+^ imaging results, we observed a significant diagnosis effect on the MFR, whereby PTHS lines showed a reduction in spontaneous network activity compared to controls (Figure 5C, D; Supp Figure 13C). Infection of PTHS lines with the RIMBP2 lentivirus resulted in a significant treatment effect, as PTHS lines showed an increase in MFR compared to PTHS control lines (Figure 5C, D). Consistent with our Ca^2+^ imaging assay, PTHS patient lines had a significant reduction in network synchronicity across development (Figure 5E, Supp Figure 13D). In addition, PTHS neurons showed reductions in burst frequency (Supp Figure 13F), and increases in the coefficient of variance for the inter-event interval (Supp Figure 13E). Of these three measures, RIMBP2 expression in PTHS neurons was effective at rescuing synchronicity (Figure 5E, Supp Figure 13G) and burst frequency (Figure 5F, Supp Figure 13H). Together, these results indicate that TCF4-mediated down regulation of RIMBP2 expression across development leads to a significant reduction in spontaneous network activity, and when RIMBP2 expression is rescued in PTHS cultures, recovery of spontaneous network activity is achieved.

Since RIMPB2 is known to regulate transmitter release within presynaptic terminals, we next tested the presynaptic specificity of this RIMBP2 rescue by performing targeted whole-cell recordings on uninfected (GFP-) neurons that are postsynaptic to RIMBP2 infected presynaptic neurons (Figure 5B). Consistent with prior results (Figure 2A-C), uninfected PTHS patient neurons in GFP alone infected cultures showed significant reductions in sEPSC frequency and amplitude compared to uninfected control neurons in GFP alone infected cultures (Figure 5G-I). However, RIMBP2 expression in presynaptic neurons resulted in a significant rescue of the sEPSC frequency in uninfected PTHS neurons (Figure 5G, H). Within diagnosis analysis revealed no effect of line on sEPSC frequency, as each line showed increased sEPSC frequency in response to RIMBP2 recovery (Supp Figure 14A, B). Importantly, upregulation of presynaptic RIMBP2 showed no treatment or within diagnosis effects on the sEPSC amplitude, further demonstrating the presynaptic specificity of the RIMBP2 rescue (Figure 5G, I; Supp Figure 14C, D). These results indicate that patient-specific mutations in *TCF4* lead to downregulation of RIMBP2 expression in presynaptic terminals, which reduces the frequency of sEPSCs and subsequently impacts spontaneous network activity. Overall, these results demonstrate TCF4 is a critical upstream regulator of synaptic development in human neurons, and we pinpoint RIMBP2 in the presynaptic terminal as a potential therapeutic target for PTHS.

## Discussion

PTHS patient neurons demonstrated robust consistency and reproducibility across a variety of phenotypes, which include delayed intrinsic excitability, network wide reductions in mean firing rates, burst frequency and synchronicity, reduced synapse density and spontaneous synaptic transmission, as well as reduced magnitude of homeostatic scaling. These phenotypes were supported by transcriptional dysregulation of genes related to neurodevelopment, presynaptic neurotransmission and synaptic plasticity. We identify *RIMBP2* as the most differentially expressed gene in PTHS patient lines and demonstrate its upregulation can rescue deficits in spontaneous network activity and presynaptic transmission. Overall, we conclude that TCF4 is an essential transcriptional regulator of excitatory neuron development and synaptic physiology, a finding which has important implications not only for PTHS, but also a number of neuropsychiatric disorders associated with TCF4.

### TCF4 as a developmental regulator of intrinsic excitability

The phenotypes identified in PTHS patient lines corroborates several previous findings from various models of PTHS. For example, PTHS patient neurons initially showed a reduction in the rate of membrane maturation which were related to intrinsic excitability deficits (Supp Figure 3). However, by week 12, these phenotypes generally resolve, suggesting they represent a delay in neuronal maturation. Similar intrinsic excitability deficits were previously reported in both human and murine models, however the developmental delay reported here appears to be a novel aspect of our *in vitro* human model. Neurons in the current study transcriptionally approximate the late third trimester to early infant postmortem human DLPFC neurons, which allows us to capture later stages of neuronal maturation not previously observed in other human models (Figure 1E). In rodent models of PTHS, two prior studies showed that TCF4 nonsense mutations resulted in similar alterations in membrane properties and intrinsic excitability in cortical and hippocampal brain slices from adolescent and adult animals ^16,22^. Moreover, in a study using hIPSC-derived neurons from a different set of PTHS patient lines than presented here, Papes and colleagues (2022) observed similar intrinsic excitability deficit in their 12wk cortical neurons to those observed in our 8wk neurons (Supp Figure 3). However, they also reported more severe phenotypes, which included a global reduction in neurogenesis and severe microcephalic-like phenotypes, which were not apparent in our 2D differentiation protocol and are not consistent with PTHS mouse models or the clinical observations of postnatal microcephaly, a clinical characteristic that is not fully penetrant, occurring in approximately 21% of patients ^13,22,47^. Furthermore, the inclusion of rodent astrocytes in our protocol, which has been found to enhance neuronal maturation of human neurons ^26,28^, may explain the larger membrane capacitance, elevated max AP output and transcriptional approximation to later stages of human cortical development that we observed in our study. Although differences in the exact developmental timing of TCF4-dependent intrinsic excitability phenotypes between human and animal models are apparent, the convergence of these phenotypes solidifies the importance of TCF4 in regulating the development of neuronal membrane properties and intrinsic excitability.

### TCF4 regulates transcription in response to neuronal activity: homeostatic plasticity

For the first time, we show Ca^2+^ imaging can be used to assay homeostatic plasticity in human neurons (Figure 4). Blocking network activity in control neurons with TTX resulted in a robust upscaling of network activity due in part to an increase in AMPA receptor accumulation and activity-dependent transcription. However, in PTHS lines, the magnitude of network upscaling and AMPA receptor accumulation was significantly reduced (Figure 4). This diagnosis effect appears to be due to regulation of activity-dependent transcription downstream of TCF4, as control lines showed a robust differential expression in genes enriched for processes related to synapses, presynaptic terminals, and synaptic plasticity, whereas no DEGs were identified from the PTHS lines following homeostatic plasticity (Supp Figure 10). TCF4’s role in activity-dependent transcription and synaptic plasticity is well documented in animal models. In response to neuronal activity, TCF4 is directly phosphorylated by PKA which is a necessary step for transcriptional activation ^20^. In addition, binding of calmodulin to TCF4, in a Ca^2+^-dependent manner, can inhibit TCF4 transcription ^48^. The convergence of these two activity-dependent signaling pathways were shown to be important for TCF4-dependent regulation of the columnar distribution of cortical neurons ^15,20^. Moreover, several PTHS mouse models show enhanced hippocampal long-term potentiation and abnormal learning and memory ^21,22^. Although homeostatic plasticity has not been previously studied in in human cells or in PTHS models, defects in synaptic scaling were previously observed in models of Fragile X and Rett’s syndrome ^49,50^ and many genes associated with neurological disorders are either required for synaptic scaling or regulated in response to induction of synaptic scaling ^40,51^.

### TCF4 regulates synaptic development and function

The most consistent phenotype we report across network development, using a variety of assays (Ca^2+^ imaging, MEA, sEPSC frequency), is a deficit in network-wide neuronal spiking. Importantly, these phenotypes are consistent across human and mouse PTHS models ^52,53^. However, we extended these findings by identifying several cellular and molecular mechanisms underlying the reduced network activity in these PTHS models. First, we showed that PTHS patient-derived neurons have a reduction in excitatory synapse density (Figure 2G-J; Figure 4E, F), a phenotype that was previously reported in the cortex and hippocampus of two different PTHS mouse models ^13,54^. Similarly, disruption of Da, the *TCF4* orthologue in drosophila, led to decreased levels of the synaptic proteins synapsin and disc large 1 ^55^. Secondly, we observed a significant reduction in the frequency and amplitude of sEPSCs, which was consistent with synaptic density measurements (Figure 2A-C; Figure 5G-I). A similar reduction in sEPSC frequency was observed in chemosensitive retrotrapezoid nucleus neurons in a PTHS mouse model ^53^. Third, mutations in *TCF4* resulted in significant dysregulation of developmental and activity-dependent gene transcription, specifically related to synaptic function, neurotransmission and synaptic plasticity (Supp Figure 10; Supp Figure 11). Differential expression analysis between PTHS and control neurons identified enrichment for genes related to the presynaptic membrane, terminal bouton, synaptic vesicle membrane, and SNARE binding, which identifies TCF4 as an important transcriptional regulator of presynaptic neurotransmitter release. This conclusion is further supported by gene set enrichment analysis of TCF4 target genes identified from ChIP-seq in SH-SY5Y cell line and differential gene expression analysis across 5 different PTHS mouse models which both identified enrichment of synaptic genes ^12,56^. All together, this convergent group of cellular and molecular mechanisms likely contribute to the robust reduction in network activity. The remarkable consistency of these phenotypes and cellular mechanisms across both human and animal models increases the likelihood they represent relevant pathophysiology in PTHS and establishes TCF4 is an important transcriptional regulator of excitatory synapse development and physiology.

### Dysregulation of RIMBP2 in PTHS patient-derived neurons

Given the consistent TCF4-dependent reduction in network activity, it was striking to find the top DEG was *RIMBP2*. It appears TCF4 may be a direct transcriptional regulator of *RIMPB2*, as *RIMBP2* expression was severely downregulated at both the NPC and neuronal stages (Figure 5A). In addition, prior ChIP-seq studies identified binding of TCF4 to the *Rimbp2*/*RIMBP2* gene locus in rat NPCs and SH-SY5Y cell line ^16,46^.

*RIMBP2* encodes a presynaptic binding protein previously shown to regulate transmitter release to varying levels, depending on the cell type and synapse. For instance, in *Drosophila*, the RIMBP2 ortholog DRBP was shown to be essential for neurotransmitter release through its coupling of synaptic vesicles (SVs), Ca^2+^ channels, and SV fusion machinery ^57^. In cultured hippocampal neurons, CA3-CA1 synapses in acute brain slices, and at the Calyx of Held synapse, RIMBP2 was shown to regulate the probability of glutamate release via its clustering of calcium channels ^43,45,58^, whereas in hippocampal mossy fiber synapses, RIMBP2 was shown to have a more substantial impact on neurotransmitter release through its interaction with Munc13-1 at presynaptic active zone; a gene also found to be significantly down-regulated in PTHS neurons (Supp Figure 11B)^43–45,57–59^. Thus, we reasoned that downregulation of *RIMBP2* expression led to a reduction in glutamate release, and subsequently disrupted the frequency of network activity in our PTHS patient neurons. Indeed, expression of RIMBP2 in PTHS cultures effectively rescued spontaneous network activity by increasing the MFR, burst frequency and synchronicity to control levels (Figure 5). Importantly, using patch-clamp to target uninfected postsynaptic neurons, we demonstrated normalization of network activity by RIMBP2 was due to presynaptic rescue of spontaneous glutamate release (Figure 5G, H), thus identifying abnormal presynaptic function as a critical factor leading to reduced network activity. These results are the first demonstration of increased RIMBP2 expression leading to enhanced function at human excitatory presynapses and are comparable with previous studies of RIMBP2 function in murine and invertebrate synapses.

In conclusion, our novel results demonstrate that TCF4 is a critical transcriptional regulator of human synaptic development, function, and plasticity. We exemplify TCF4’s role in modulating synapse development and function via its transcriptional regulation of *RIMBP2* and established that dysregulation of RIMBP2 in presynaptic terminals as a molecular mechanism underlying abnormal synapse development and network excitability in PTHS. Furthermore, these findings highlight RIMBP2 as a potential therapeutic target for recovery in the broader patient population associated with TCF4-specific dysfunction.

## Materials and Methods

### Generation and maintenance of hiPSCs

Early passage fibroblasts (less than passage 5) were thawed and cultured in cDMEM media (DMEM, 10% Fetal Bovine Serum, 1% Non-Essential Amino Acids (NEAA), beta-mercaptoethanol (b-ME) and 1% penicillin/strep). According to the manufacturer’s protocol ^60^, all cell lines were reprogrammed using Cytotune™-iPS 2.0 Sendai Reprogramming Kit (A16517; ThermoFisher Scientific). Clones obtained were cultured onto irradiated mouse embryonic fibroblast (ir-MEF) feeder layer in hESC media (DMEM/F-12, 20% Knockout serum replacement (KSR), 1% NEAA, b-ME) supplemented with FGF2 (10ng/ml; 100-18B; Peprotech) to prevent spontaneous differentiation of iPS colonies with daily medium changes. iPS cells were passaged once a week (∼80% confluence/well) at 1:4-1:6 ratios using 1mg/ml collagenase (Gibco) to detach colonies and onto fresh ir-MEF plates.

### Validation of hiPSC pluripotency

For stem cell quality control, immunocytochemistry of pluripotency markers (NANOG, SOX2, OCT3/4 and TRA 1-60), quantitative PCR for pluripotency markers (NANOG, OCT3/4, SOX2, KLF4 and LIN28), Karyotyping (G-banding) was performed on all iPS cell lines used in the study (Supp Figure 1A). For pluripotency marker staining (see ICC methods), primary antibodies against NANOG, SOX2, OCT3/4 and TRA 1-60 were diluted in the blocking buffer and incubated overnight at 4? (Supp Figure 1A), only lines with contiguous high expression across all markers were banked and used in this study. For quantitative PCR (qPCR), total RNA was extracted from iPS cells (see RNA extract methods below) and analyzed for expression of pluripotency markers. The relative fold changes in expression were calculated using the 2-ΔΔCt method relative to GAPDH and fibroblasts as internal controls. For karyotyping analysis, cells were grown on 25cm^2^ tissue culture flasks and karyotyping was performed at WiCell (chromosome analysis service). Sendai virus staining (PD029; MBL) was performed to confirm clearance of Sendai Virus in the hiPS clones.

### Differentiation into cortical neurons

hiPS cells were differentiated into cortical neurons as previously described in (Wen et al, 2014), with modifications. Briefly, hiPSC colonies were detached from the ir-MEF feeder layer by incubating the cells with collagenase, type IV (1mg/ml; Gibco) for 1h at 37?. The detached colonies were cultured in suspension in ultra-low attachment 10 cm culture dishes (Corning) in FGF-2 free hESC media supplemented with Dorsomorphin (2µM) and A-83 (2µM) (here on referred to as embryoid body (EB) media) for 4 days with daily medium changes. On day 5, EB media was replaced with neural induction (hNPC) media consisting of DMEM/F-12, N2 supplement and NEAA supplemented with heparin (2mg/ml) and cyclopamine (2µM). On day 7, EBs growing in suspension were then transferred onto Matrigel-coated 10 cm dishes to form neural tube-like rosettes. The attached rosettes were cultured for 10 days with media changes every other day, after 10 days, rosettes were mechanically picked using hand-pulled glass pasteur pipettes and transferred to ultra-low attachment 6 well plates. Picked rosettes were cultured for additional 3-4 days in hNPC medium containing B27 forming neural spheres. For neuronal differentiation, neural spheres were dissociated with Accutase (Gibco) at 37°C for 10 min. Cells were passed through a 40µm mesh filter to obtain a single cell suspension of neural progenitor cells (NPCs). NPCs were then plated at 150,000 cells per well, on pre-treated poly-L-ornithine/laminin plates with a confluent layer of primary rat astrocytes (see below). NPCs were plated on either 24-well ibidi for fate ICC and Ca^2+^ imaging; pretreated glass coverslips for synaptic ICC and electrophysiology; standard 24-well plates for biochemical analysis, or 48-well Axion Cytoview MEA plates (Catalog number: M768-tMEA-48B). Neuronal cultures were treated DAPT (10µM) at DIV21-28 days in culture; and mitotic arrest was induced at DIV28 via AraC (4µM). Neuronal cultures were maintained for up to 18wks with half medium changes twice a week with Neuron medium (Neurobasal Plus media, 1% Glutamax, B27^+^, supplemented with Ascorbic Acid (200nM), cAMP (1µM), BDNF (10mg/ml) and GDF (10mg/ml)).

### Primary rat astrocytes

Primary astrocytes were prepared as previously described with modifications ^61,62^. Briefly, cortices were extracted, and meninges removed from postnatal day 2–3 pups. After dissection cortices were triturated with a 1ml pipette 10 times to break down the tissue into smaller pieces. The tissue was then incubated with 300µl 2.5% trypsin-EDTA (Gibco) and 100µl DNase (100µg/ml; Sigma) on a shaker at 150 rpm at 37? for 30-45 minutes. Subsequently, the cells were passed through a 40µm mesh filter, and transferred in Glial media (MEM, 5% FBS, Glucose (20mM), Sodium bicarbonate (0.2mg/ml), L-Glutamine (2.5mM) and 1% Pen/Strep) to poly-L-ornithine/laminin coated 75cm^2^ tissue culture flasks (5-6 cortices per flask). After 24h, the flasks were vigorously tapped and shaken to remove any loosely attached cells. The cells were cultured for 5-7 days with medium changes every other day, until ∼80% confluence, after which they were shaken at 150 rpm in 37? overnight. The next day cells were passaged at a 1:3 ratio before final passage onto poly-L-ornithine/laminin pre-coated 24-well plates (initial seeding density 70,000 cells/well). Plated astrocytes were mitotically arrested via cytosine arabinoside treatment (AraC; 20µM) once wells were ∼80% confluent and maintained with astrocyte media until NPC plating < 1wk after.

### Immunocytochemistry

Cells were fixed in phosphate-buffered saline (PBS) containing 4% paraformaldehyde (PFA) for 15 min at room temperature (RT). Cells were blocked with 10% Goat serum and 0.3% Triton X-100 in PBS for 1hr at RT; followed by overnight incubation with primary antibodies in fresh blocking solution. For all surface marker ICC staining, detergent concentration was reduced to 0.15% Triton X-100. The cells were then rinsed with PBS and incubated with species-specific Alexa Fluor 488-, Alexa Fluor 555-, Alexa Fluor 568-, or Alexa Fluor 647-conjugated secondary antibodies (1/1,000; Thermo Fisher) in PBS with 0.15% Triton X-100 for 90 mins at RT, followed by staining with DAPI (4′,6-diamidino-2-phenylindole) for 5 min to capture the nuclei. All images were acquired using a Zeiss 780 confocal microscope. The details of primary and secondary antibodies are described in Supp Table 9.

### RNA isolation and qPCR

Total RNA was isolated from samples using phenol:chloroform isolation with Trizol followed by purification using the RNeasy Micro Prep kit (Qiagen), and quantified using a Nanodrop. Subsequently, cDNA was prepared using High-Capacity cDNA Reverse Transcription Kit (4387406; Thermofisher Scientific). qPCR assays were carried out in duplicate or triplicate using the QuantStudio3 Real-Time PCR system (Applied Biosystems). qPCR was performed using Taqman probes (Supp Table 9) according to manufacturer’s instructions.

### RNA sequencing data generation and processing

RNA was collected from three different developmental stages, NPCs (DIV0), Neurons at 8wks and 18wks; and alongside 8wk baseline, 8wk cultures treated with TTX for 48hr. Illumina RNA sequencing (RNA-seq) libraries were constructed from RNA with Ribo-Zero gold kits across three batches and included ERCC spike-in sequences. Libraries were prepared and sequenced on an Illumina HiSeq 3000 at the Lieber Institute. RNA-seq data read alignment and quantification was performed using a custom human ^+^ rat annotation as previously described ^26^. Gene expression counts from Gencode v25-annotated genes were considered for all subsequent analyses (and rodent gene counts were excluded from analysis).

### Transcriptional analysis

After quality control checks, principal component analysis (PCA) was performed on log2-transformed gene expression levels (using the RPKM summarization) across the three *in vitro* developmental stages generated in this manuscript and compared with that of human Dorsolateral Prefrontal Cortex (DLPFC) postmortem samples, from prenatal stages through later adulthood. We performed RNA deconvolution using human neural cell types as previously described to quantify neuronal maturity ^26^. Differential gene expression by diagnosis, time or treatment was then performed using the limma voom approach ^63^, adjusting for the first PC of RNA deconvolution proportions, an approximation of the the human neuron plating proportion (calculated as the fraction of total human ^+^ rat reads that were from human reads) and the human gene assignment rate, which is the fraction of human-aligned reads that were annotated to exonic sequences. Finally, Gene Ontology analysis was conducted on the DEGs enriched in NPC and neuron cultures (8wk and 18wk samples) and after homeostatic plasticity using the clusterProfile tool ^64^.

### Whole-cell patch-clamp recordings

Whole-cell electrophysiology recordings were performed as previously described using the following solutions ^26^. Extracellular recording buffer contained (in mM): 128 NaCl, 30 glucose, 25 HEPES, 5 KCl, 2 CaCl_2_, and 1 MgCl_2_ (pH 7.3). For current- and voltage-clamp measurements, pipettes were filled with (in mM): 135 KGluconate, 10 Tris-phosphocreatine, 10 HEPES, 5 EGTA, 4 MgATP, and 0.5 Na_2_GTP (pH 7.3). Patch pipettes were fabricated from borosilicate glass (N51A, King Precision Glass, Inc.) to a resistance of 2-5 MΩ. Current signals were recorded with either an Axopatch 200B (Molecular Devices) or a Multiclamp 700A amplifier (Molecular Devices) and were filtered at 2 kHz using a built in Bessel filter and digitized at 10 kHz. Voltage signals were filtered at 2kHz and digitized at 10kHz. Data was acquired and analyzed using Axograph on a Dell PC (Windows 7). For voltage clamp recordings, cells were held at -70 mV for sodium/potassium currents and sEPSCs, and 0 mV for sIPSCs. After recording a ramp protocol (−110 to 110 mV, 0.22 mV/s) for each neuron in voltage clamp, the recording configuration was switched to current clamp and a series of APs (20 episodes) was elicited at 0.5 Hz with a square current injection that was increased by 10 pA every episode. Calculation of dV/dT maximum and minimum was set as the maximum slope for the ascending and descending phases with three regression points per maximum slope.

### Calcium Imaging and analysis

Neurons were infected at ∼5wks with AAV1-hSyn1-mRuby2-P2A-GCaMP6s (a gift from Tobias Bonhoeffer, Mark Huebener and Tobias Rose (Addgene viral prep # 50942-AAV1; http://n2t.net/addgene:50942 ; RRID:Addgene_50942) ^65^. Imaging was performed in culture media biweekly from 6wks to 18wks in culture on a Zeiss LSM780 equipped with a 10X/0.45NA objective and a temperature- and atmospheric-controlled enclosure. A reference image was acquired for each field of mRuby fluorescence, then a time-series was acquired at 4Hz for 8 minutes. Regions of interest were identified from the red reference image, and fluorescent intensity over time was computed for each ROI then normalized to ΔF/F. Peaks were identified and the number of events was calculated for each ROI. The Ca^2+^ synchronicity index incorporates both per-field and individual per-ROI neuronal activity metrics. Correlation maps were constructed to compare the pattern of fluorescence intensity changes with known motifs representing calcium events, using 16 previously published motifs, as well as 7 new motifs representative of Ca^2+^ activity in these cultures ^66^. Synchronicity index per field was then calculated and compared across baseline and treatment conditions (e.g. 48hrs TTX (1µM). Acute application of PTX (100µM) was applied in 8wk cultures, at a stage when sIPSCs were significantly different between PTHS and control cultures. Acute application of dl-AP5 (100µM) ^+^ DNQX (10µM) was applied at the end of the experiment, in 18wk cultures, when cultures were most active.

### Homeostatic plasticity

In DIV54 cultures 2 wells per line previously infected with AAV1-hSyn1-mRuby2_P2A_GCaMP6s virus were treated with 1µM TTX. After 24hrs 1 treated well was aspirated, PBS was used to rinse the well 1X, before 250µl of conditioned media from a neighboring non-treated well of the same line was used to replace TTX treated media. Freshly rinsed wells and control wells were then imaged for Ca^2+^ activity according to the protocol above, within ∼15 minutes of media replacement. After 48hrs the remaining treated well per line was aspirated, PBS was used to rinse the well 1X before 250ul of conditioned media from a neighboring non-treated well was used to replace TTX treated media. Again, freshly rinsed wells and control wells were then imaged for Ca^2+^ activity according to the protocol above, within ∼15 minutes of media replacement.

### Multi-Electrode Array (MEA) preparation and analysis

Cytoview MEA plates (M768-tMEA-48B) were prepared with initial surface treatment using 70% EtOH for 10 minutes at room temperature (RT). The wells were then pre-treated with poly-L-ornithine/laminin, each incubated at RT for 1h followed by 2 washes with ultraPure water. The wells were allowed to dry and replaced with HBSS prior to plating. A 10uL/well cell suspension of dissociated NPCs (as described above) at a density of 150,000 cells/well and primary rat astrocytes (as described above) at a density of 70,000 cells/well was prepared and plated to cover the electrode in each well. The cultures were incubated at 37°C, 5% CO_2_ for 1hr for the cells to settle down on the electrodes, after which an additional 300-400uL of Neuron media was added to each well. The neuronal cultures were maintained with half media changes twice a week as described above. MEA recordings were performed using the Axion Meastro MEA system (Axion Biosystems). The plates were recorded on stage heated to 37°C and ventilated with a mixture of 5% CO_2_:95% air. Recordings were performed in Axion Integrated Studio (AxIS) Navigator (Axion Biosystems) software. The plate was first allowed to equilibrate for 5 minutes in the Maestro device, and then activity was recorded for 10 minutes. Each plate was recorded for 10 mins, once a week from 6wks to 12wks. Spikes were detected with AxIS software using an adaptive threshold crossing set to 5.5 times the standard deviation of the estimated noise for each electrode. Raw data was analyzed using the Axion Biosystems Neural Metrics Tool software (version 3.1.7). Electrodes that detected at least 5 spikes/min were classified as ‘active electrodes’ for the analysis. Bursts were identified from each individual electrode using an inter-spike interval (ISI) threshold requiring a minimum number of 5 spikes with a maximum ISI of 100 ms. A minimum of 50 spikes with similar maximum ISI of 100 ms with a minimum of 35% active electrodes were required for network bursts in the well. The synchrony index was calculated using a cross-correlogram synchrony window of 20ms. Recordings took place 24hrs after the previous feeding.

### Statistics

All values graphed are expressed as mean ± s.e.m. Statistical significance for ICC, electrophysiology, qPCR and MEA metrics were calculated using GraphPad Prism (GraphPad Software) and consisted of two-tailed t-tests for groups of two, one-way ANOVAs when groups were >2 and two-way ANOVAs when multiple conditions are compared, with Tukey’s or Šídák’s multiple comparisons test values reported for follow up tests and multiple test correction. For Ca^2+^ events, statistics were calculated using MATLAB (MathWorks) using a mixed model. For transcriptional analysis R statistical programming language (R Core Team, 2017) was used.

## Acknowledgements

We are grateful for the vision and generosity of the Lieber and Maltz families, who made this work possible. We thank Dr. Christian Rosenmund’s lab for sharing the *Rimbp2* lentiviral construct and Dr. Stephen J. Haggarty’s lab, Dr. David Sweetser’s lab, and Dr. Guo-li Ming’s lab for sharing PTHS patient cell lines. This work was supported by the Lieber Institute for Brain Development, the Pitt-Hopkins Research Foundation Awards (to B.J.M.), National Institute of Mental Health (NIMH) grant R01MH110487 (to B.J.M.), R01MH097949 (to B.W.L.), and NIMH training grant T32MH015330 (to B.A.D.). The content is solely the responsibility of the authors and does not necessarily represent the official views of the National Institutes of Health.

## Author Contribution

Conceptualization, B.A.D., and B.J.M.; methodology, B.A.D., M.T., A.E.J., and B.J.M.; formal analysis, B.A.D., H.Y.C., Z.Y., I.O., F.F., M.T., and A.E.J.; resources, S.R.S., Y.H.W., R.L.M, M.L., and B.W.L; data curation, B.A.D., H.Y.C., Z.Y., I.O., S.R.S., N.M.P., D.D., J.M.M., J.F.B and G.S.; figure generation, B.A.D., M.T., A.E.J. writing – original draft, B.A.D and B.J.M.; writing – review & editing, B.A.D., D.D., I.O., S.R.S., A.E.J., and B.J.M; supervision, B.A.D., S.R.S., A.E.J., and B.J.M.; project management, B.A.D., S.R.S., and B.J.M.; funding acquisition, B.A.D., and B.J.M.

---

**Supplemental Figure 1.**
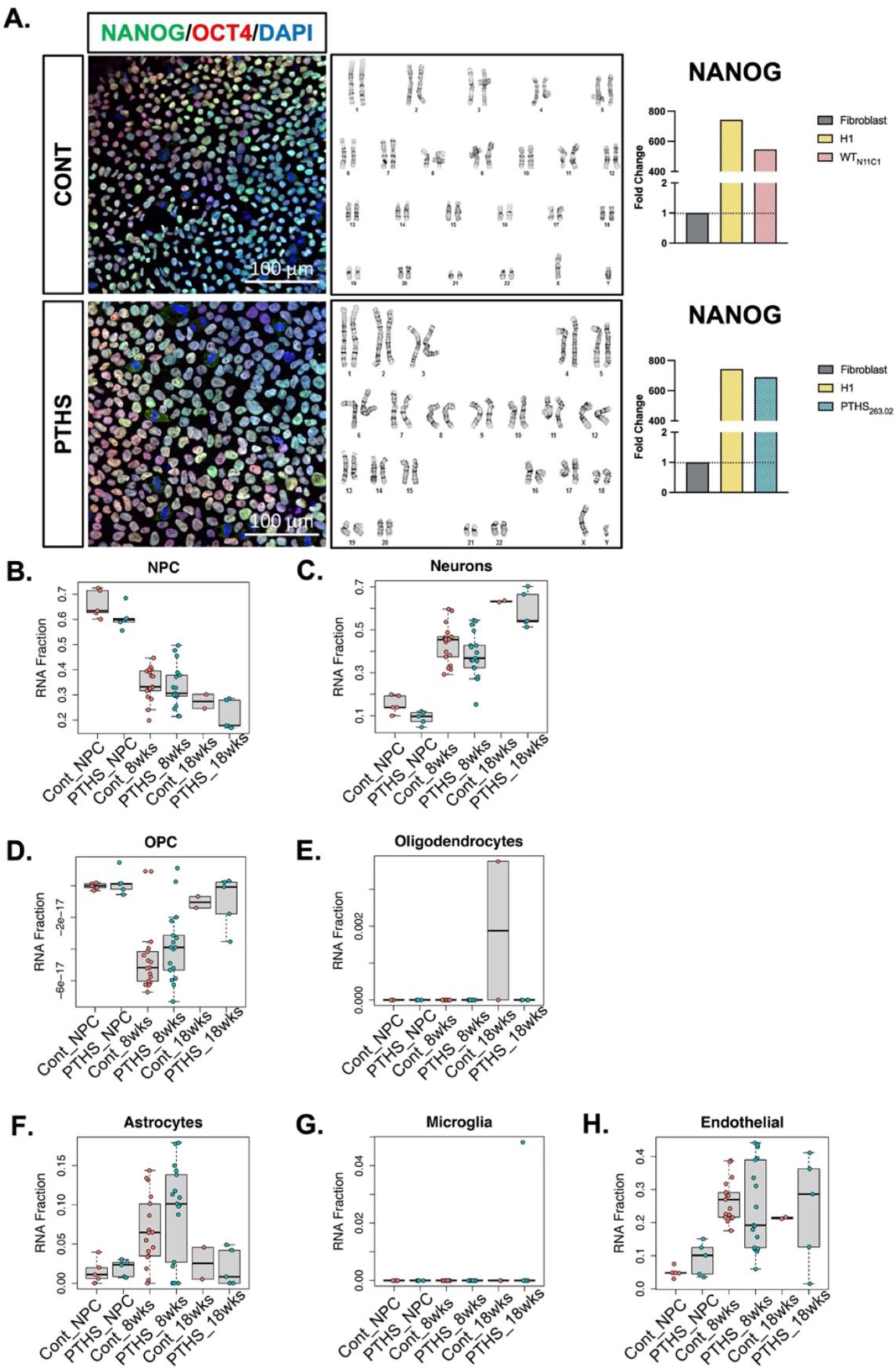
Diagnosis had no effect on hiPSC characterization or cell-type specification across development. **A**. Representative example of hiPSC characterization and quality control showing ICC staining of the pluripotency markers NANOG and OCT4, karyotyping analysis, and qPCR quantification of NANOG. **B-H** Cellular deconvolution analysis of RNA sequencing data obtained across cortical differentiation showing expected developmental changes in cellular populations which are not different by diagnosis. The fraction of RNA assigned to each cell type is plotted. Neural progenitor cells (NPCs), oligodendrocyte progenitor cells (OPCs).

**Supplemental Figure 2.**
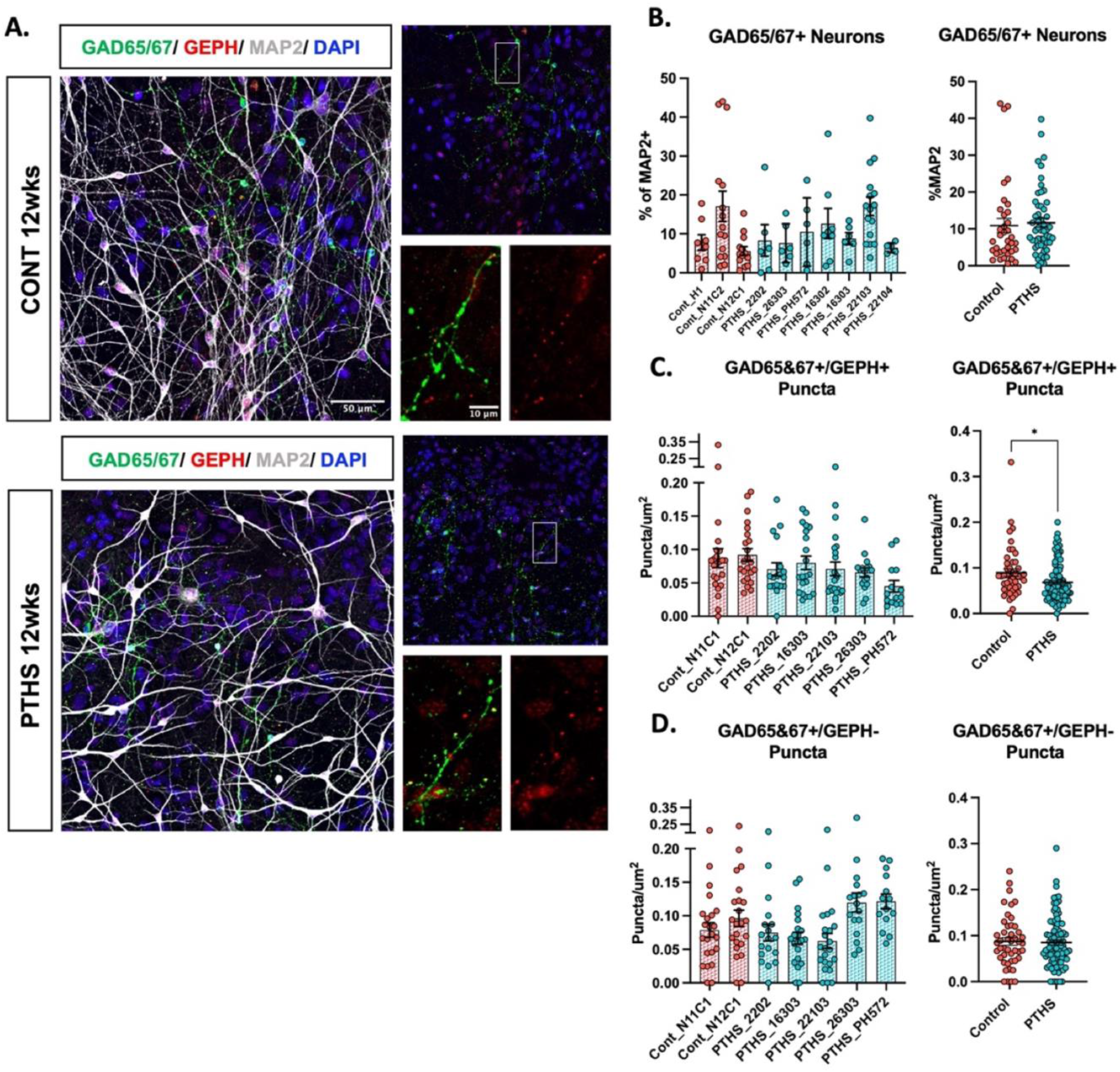
Diagnosis had no effect on the proportion of GABAergic neurons but was associated with reduced inhibitory synapses in PTHS patient-derived cortical neurons. **A**Example images of 12wk GABAergic neurons from control and PTHS cultures stained for GAD65/67 (green), gephyrin (GEPH, red) and MAP2 (white). **B**. Quantification of ICC images showing the percent of GAD65/67+ cells within the MAP2+neuron population was not different by line or diagnosis. Each data point represents the percentage of MAP2+cells co-labelled with GAD65/67, averaged across three fields per experiment; across 3 independent differentiations. **C**. Quantification of the density of GAD65/67+ GEPH+ puncta along neurites by line and by diagnosis. PTHS neurons showed a lower density of GAD65/67+ GEPH+ puncta compared to controls (*t* (1,137) = 2.505; p = 0.0134). **D**. Quantification of the density of GAD65/67+ GEPH-puncta showing no difference by line or diagnosis. Each data point in **C**. and **D**. represents the average of three independent neurite segments, which in sum are >300µm in length, averaged per field; three fields, per line, per independent experiment, across a minimum of four independent

**Supplemental Figure 3.**
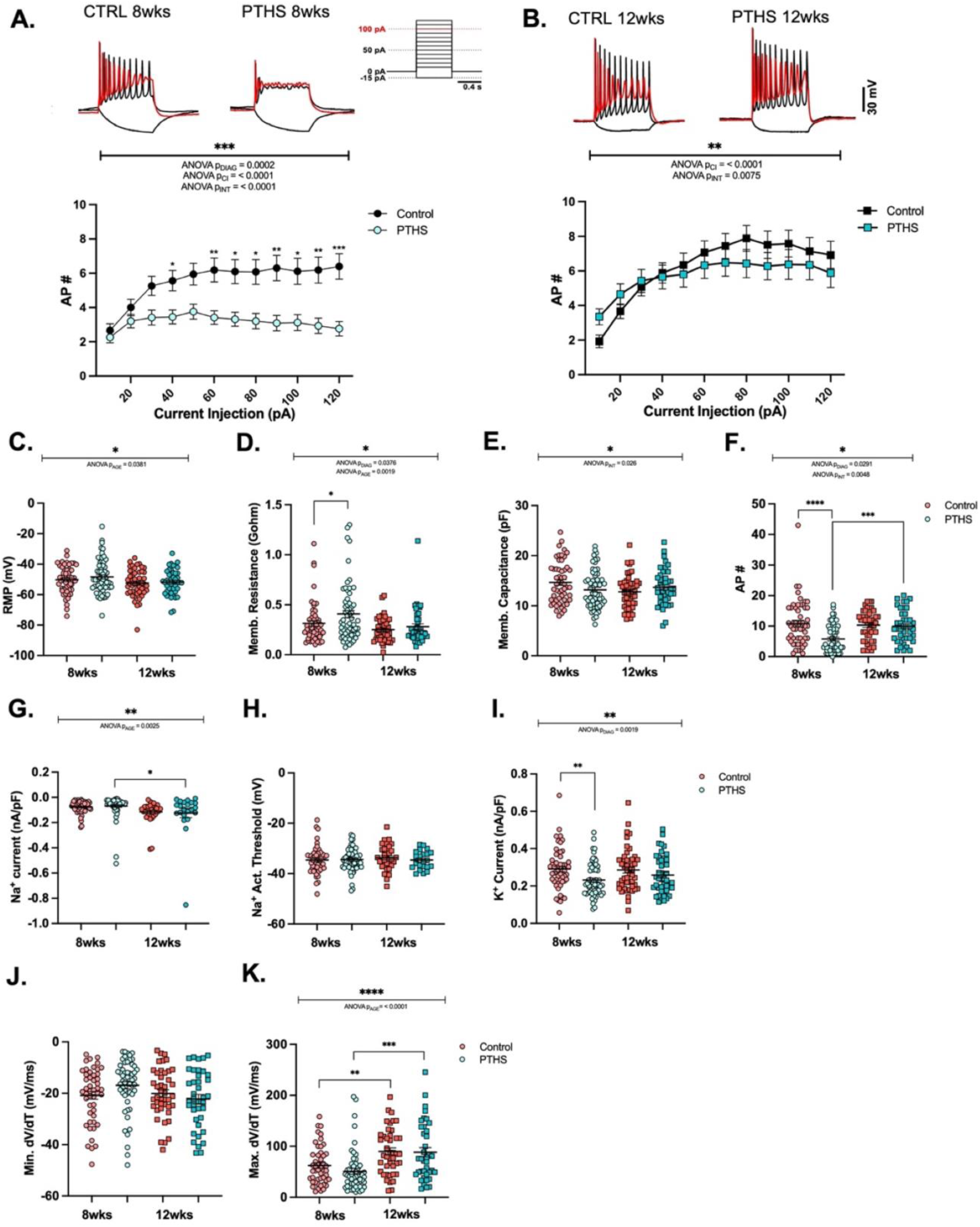
PTHS patient-derived neurons displayed developmental delays in their intrinsic excitability. **A**. Example current-clamp traces from 8wk neurons in response to 600ms current injections (+100pA red trace and -50pA black trace). Below, AP input-output curve in 8wk neurons show a main effect of diagnosis (F (1, 109) = 14.63, p *=* 0.0002), a main effect of current injection (F (2.445, 266.5) = 8.844, *p* < 0.0001), and an interaction effect between current injection and genotype (F (11, 1199) = 4.677, *p* < 0.0001). **B**. Example current-clamp traces from 12wk neurons in response to 600ms current injections (+100pA red trace and -50pA black trace). Below, AP input-output curves in 12wk neurons show a main effect of current injection (F (2.038, 165.1) = 20.75, p < 0.0001), and an interaction effect between current injection and diagnosis (F (11, 891) = 2.347, p = 0.0075). **C**. Quantification of resting membrane potential demonstrates no effect of genotype, but a main effect of age F(1,203) = 4.359, p = 0.0381). **D**. Quantification of membrane resistance demonstrates a significant increase in PTHS neurons, with a main effect of diagnosis (F (1, 203) = 4.380, p = 0.0376) and age (F (1, 203) = 9.943, p = 0.0019); with effects being driven by a significant increase at 8wks in PTHS (*t* (1,203) = 2.362, p = 0.0379). **E**. Quantification of membrane capacitance demonstrates no main effect of diagnosis, but an interaction effect of genotype and age (F (1, 203) = 5.055, p = 0.026). **F**. Quantification of maximum AP output demonstrates a significant reduction in PTHS with a main effect of diagnosis (F (1,190) = 4.846, p = 0.0291) and interaction between age and diagnosis (F (1,190) = 11.18, p = 0.0048), driven by reduced AP firing in PTHS specifically in 8wk neurons (*t* (1,109) = 4.405, p < 0.0001). **G**. Quantification of Na^+^ current density demonstrates age-dependent increases (F (1, 165) = 9.430; p = 0.0025). **H**. Quantification of Na^+^ channel activation threshold demonstrates no differences between PTHS and control neurons. **I**. Quantification of K^+^ currents density shows a main effect of diagnosis (F (1, 200) = 8.647, p = 0.0019), which was predominantly driven by differences at the 8wk time point (*t* (1,200) = 3.031, p = 0.0055). **J**. Quantification of minimal dV/dT (AP downslope) demonstrates no differences between PTHS and control neurons. **K**. Quantification of maximum dV/dT (AP upslope) demonstrates no differences between PTHS and control; however there was a significant increase with age (F(1,180) = 23.64, p < 0.0001). Each data point in **C**-**K**. represents individual neurons from five control lines and four PTHS lines, from a minimum of four independent differentiations, collapsed by diagnosis, see Supp Figure 4-6 for individual line data.

**Supplemental Figure 4.**
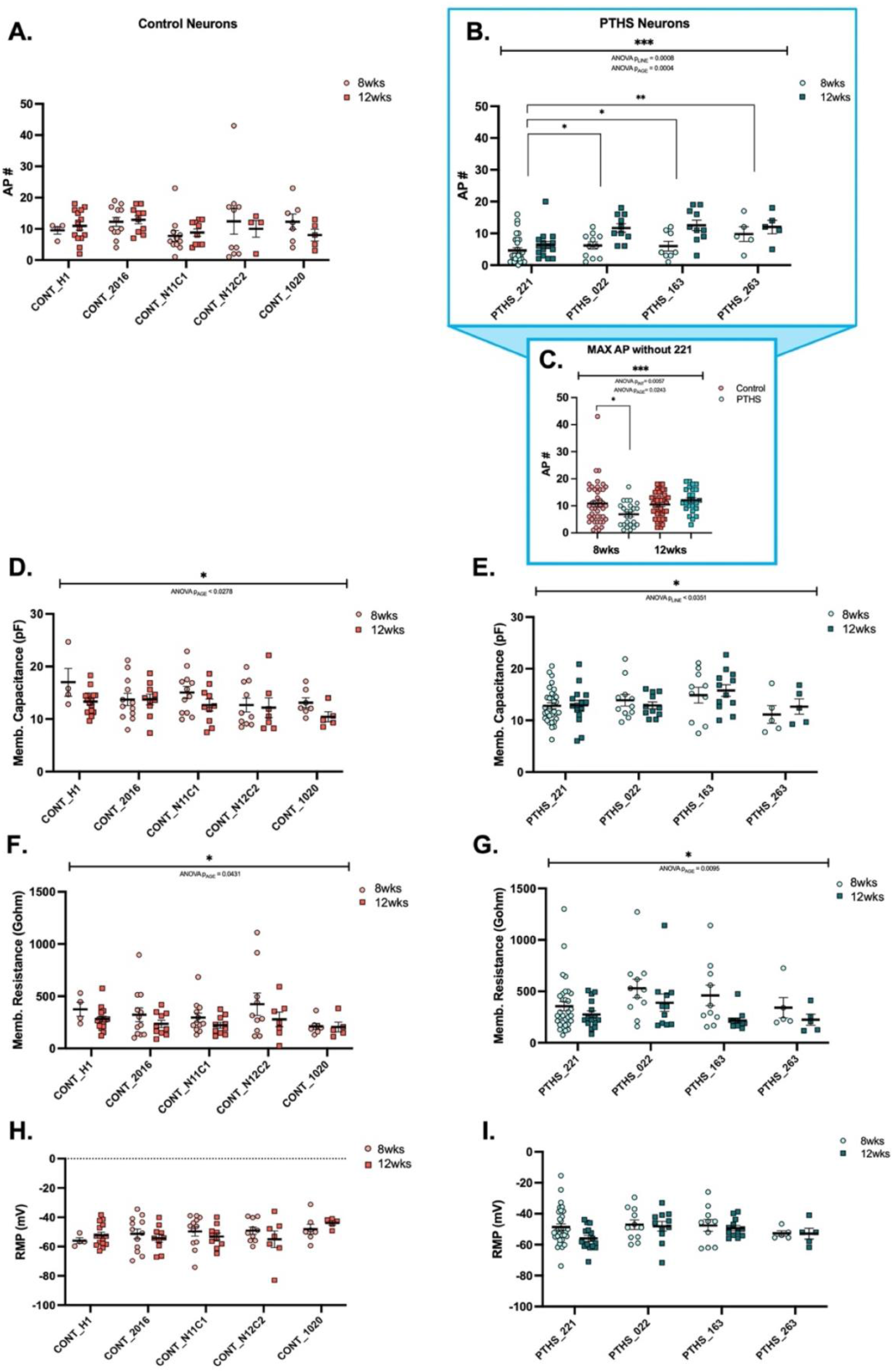
Analysis of the within diagnosis variation of intrinsic excitability. **A**. Maximum AP output was not different across control lines. **B**. Maximum AP output was significantly different within PTHS lines (F (3, 85) = 6.174, p = 0.0008); post hoc analysis demonstrates the difference is driven by PTHS patient 221: PTHS_221 vs. PTHS_022 (*q* (3,85) = 4.037, p = 0.0272; PTHS_221 vs. PTHS_163 (*q* (3,85) = 4.163, p = 0.0214; PTHS_221 vs. PTHS_263 (*q* 3,85) = 4.834, p = 0.0053). There was also a significant main effect of age within PTHS lines (F (1,85) = 13.80, p = 0.0004). **C**. Summary data showing a significant interaction effect remains in the maximum AP output after PTHS patient 221 is removed (F (1, 141) = 5.185, p = 0.0057); post hoc analysis demonstrates this effect is driven by the 8wk time point (*t* (1,141) = 2.788, p = 0.012). **D**. Membrane capacitance was not significantly different between control lines, but a significant effect of age was observed (F (1, 82) = 5.018, p = 0.0278) **E**. Membrane capacitance was significantly different between PTHS patient lines F (3.93) = 2.987, p = 0.0351) **F**. Membrane resistance was not significantly different between control lines, but a significant effect of age was observed (F (1, 82) = 4.222, p = 0.0431). **G**. Membrane resistance was not significantly different between PTHS patient lines, but a significant effect of age was observed (F (1, 93) = 7.016, p = 0.0095). **H**. Resting membrane potential was not significantly different between control lines. **I**. Resting membrane potential was not significantly different between PTHS patient lines. Each data point in **A, B, D-I**. represent individual neurons from five control lines and four PTHS lines, from a minimum of four independent differentiations. Each data point in **C**. represents line data collapsed by diagnosis.

**Supplemental Figure 5.**
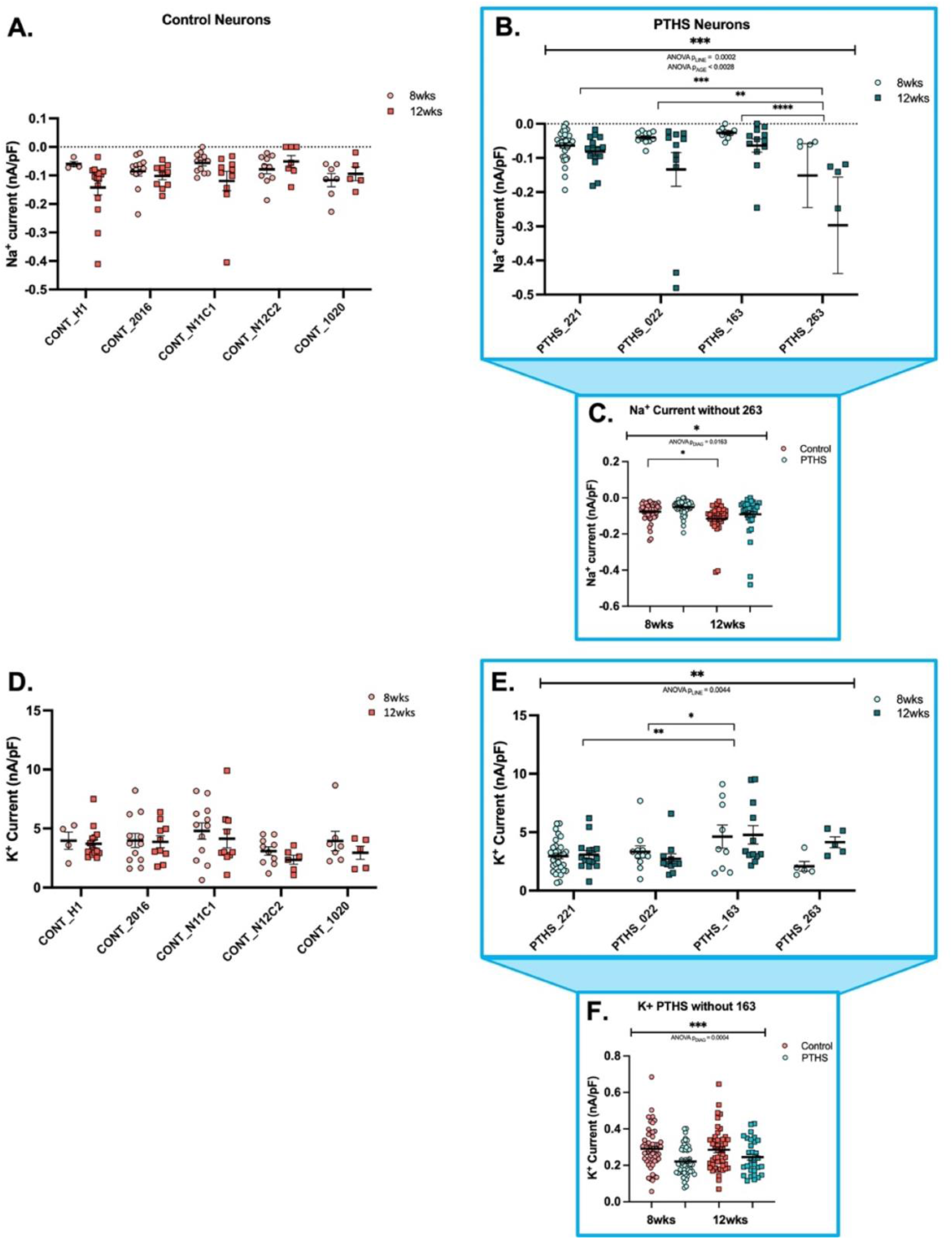
Analysis of the within diagnosis variation of voltage-gated ion channel current densities. **A**. Na^+^ current density was not significantly different between control lines. **B**. Na^+^ current density was significantly different between PTHS patient lines (F (3, 93) = 7.437, p = 0.0002), which was driven by PTHS patient line 263: (PTHS_263 vs. PTHS_022 (*q* (3,22) = 4.963, p = 0.0038); (PTHS_263 vs. PTHS_163 (*q* (3,22) = 6.462, p < 0.0001); (PTHS_263 vs. PTHS_221 (*q* (3,47) = 5.928, p = 0.0004). **C**. Summary data showing a significant diagnosis effect on Na^+^ current density after PTHS patient line 263 is removed (F (1, 173) = 5.887, p = 0.0163). **D**. K^+^ current density was not significantly different between control lines. **E**. K^+^ current density was significantly different between PTHS patient lines (F (3, 91) = 4.677, p = 0.0044) which was driven by PTHS patient line 163: (PTHS_163 vs. PTHS_221 (*q* (3,46) = 4.923, p = 0.0042); (PTHS_163 vs. PTHS_022 (*q* (3,22) = 4.340; p = 0.0148). **F**. Summary data showing a significant diagnosis effect on K^+^ current density after PTHS patient line 163 is removed (F (1, 179) = 12.98, p = 0.004). Each data point in **A, B, D, E**. represents individual neurons from five control lines and four PTHS lines, from a minimum of four independent differentiations. Each data point in **C**. and **F**. represents line data collapsed by diagnosis.

**Supplemental Figure 6.**
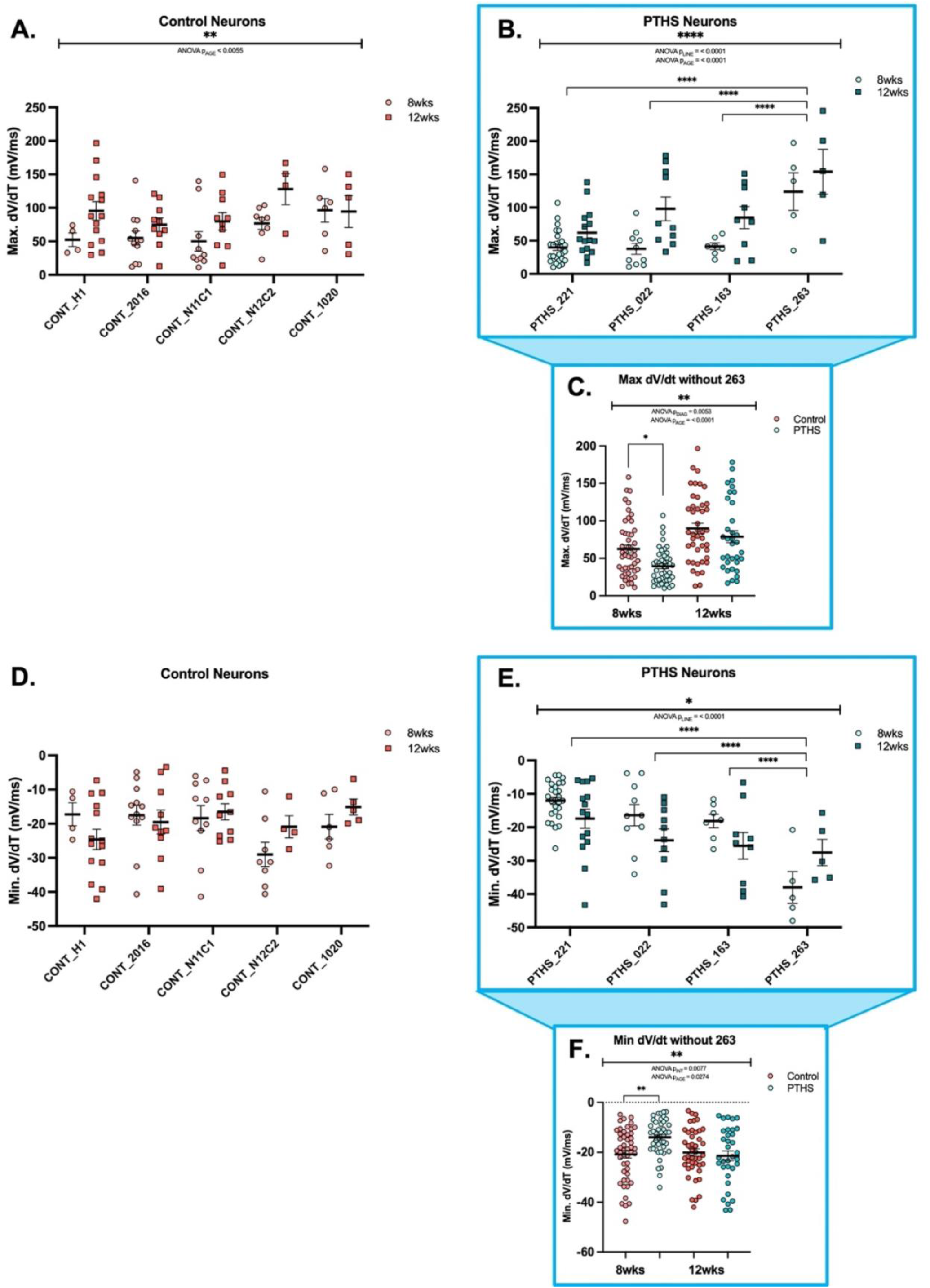
Analysis of within diagnosis variation for AP properties. **A**. Maximum dV/dT was not significantly between control lines but showed an effect of time (F (1, 73) = 8.183, p = 0.0055). **B**. Maximum dV/dT was significantly different between PTHS patient lines (F (3, 80) = 13.02, p < 0.0001), which was driven by PTHS patient line 263: (PTHS_263 vs. PTHS_221 (*q* (3,80) = 8.808; p < 0.0001); (PTHS_263 vs. PTHS_022 (*q* (3,80) = 6.522; p < 0.0001); (PTHS_263 vs. PTHS_163 (*q* (3,80) = 6.664; p < 0.0001). **C**. Summary data showing a significant effect of diagnosis after PTHS patient line 263 is removed (F (1, 165) = 7.979, p = 0.0053); post hoc analysis demonstrates the interaction was driven by the 8wk time point (*t* (1,165) = 2.81; p = 0.0111); the main effect of age remained (F (1, 165) = 30.66, p = < 0.0001). **D**. Minimum dV/dT was not significantly different between control lines. **E**. Minimum dV/dT was significantly different between PTHS patient lines (F (3, 80) = 11.12, p < 0.0001), which was driven by PTHS patient line 263: (PTHS_263 vs. PTHS_221 (*q* (1,80) = 7.946, p < 0.0001); (PTHS_263 vs. PTHS_022 (*q* (1,80) = 5.099, p = 0.003); (PTHS_263 vs. PTHS_163 (*q* (1,80) = 4.232, p = 0.0189). **F**. Summary data showing a significant interaction between diagnosis and age after PTHS patient line 263 is removed (F (1, 165) = 7.270, p = 0.0077); post hoc analysis demonstrates the interaction was driven by the 8wk time point (*t* (1,165) = 3.334, p = 0.0021); there was also a main effect of age (F (1, 165) = 4.951, p = 0.0274). Each data point in **A, B, D, E**. represents individual neurons from five control lines and four PTHS lines, from a minimum of four independent differentiations. Each data point in **C, F**. represents line data collapsed by diagnosis.

**Supplemental Figure 7.**
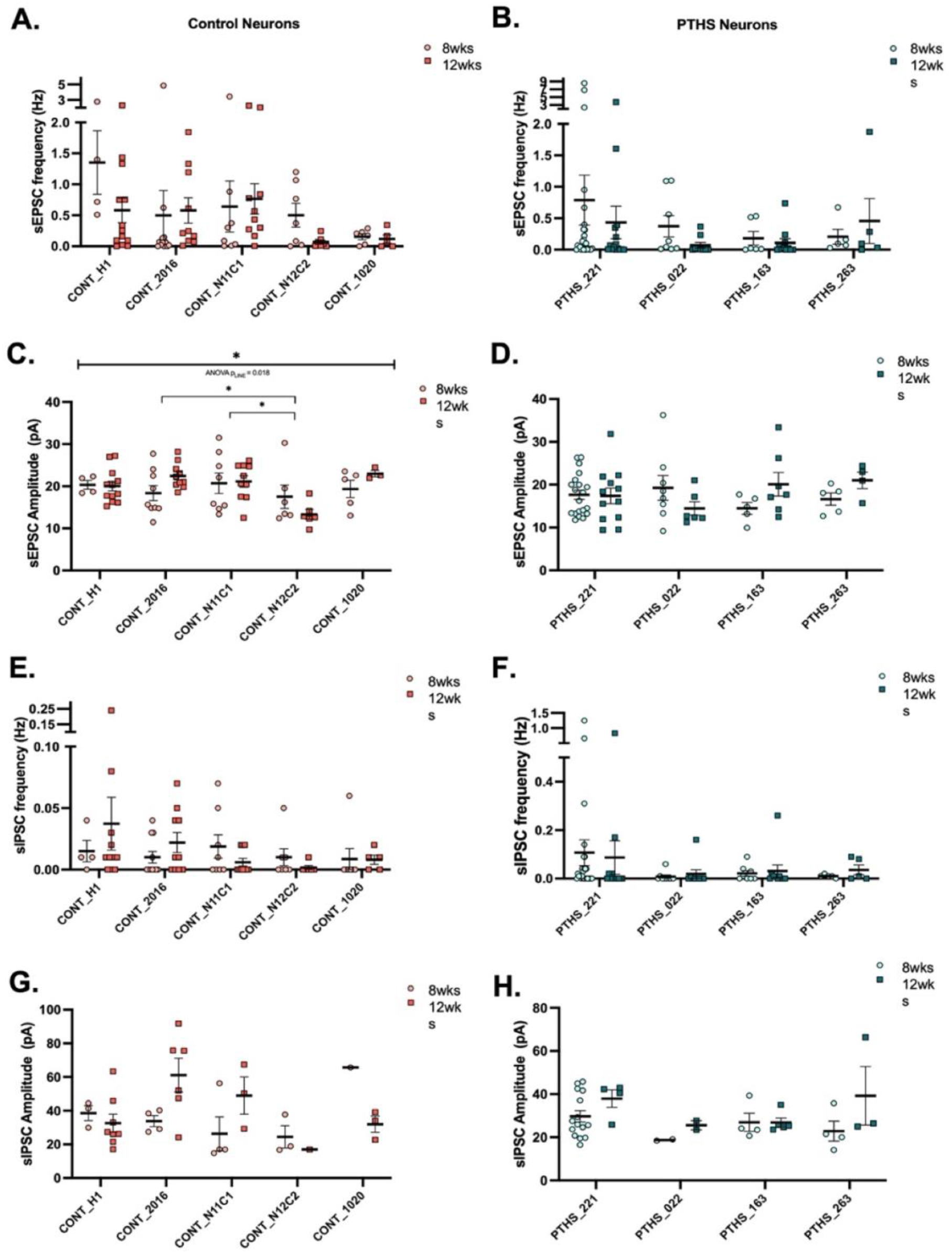
Analysis of within diagnosis variation of spontaneous synaptic transmission and synapse density. **A**. sEPSC frequency was not different between control lines. **B**. sEPSC frequency was not different within PTHS patient lines. **C**. sEPSC amplitude was significantly different between control lines (F (4, 63) = 3.195, p = 0.0188). **D**. sEPSC amplitude was not different between PTHS patient lines. **E**. sIPSC frequency was not different between control lines. **F**. sIPSC frequency was not different between PTHS patient lines. **G**. sIPSC amplitude was not different between control lines. **H**. sIPSC amplitude was not different between PTHS patient lines. Each data point in **A-H**. represents individual neurons from five control lines and four PTHS lines, across a minimum of four independent differentiations.

**Supplemental Figure 8.**
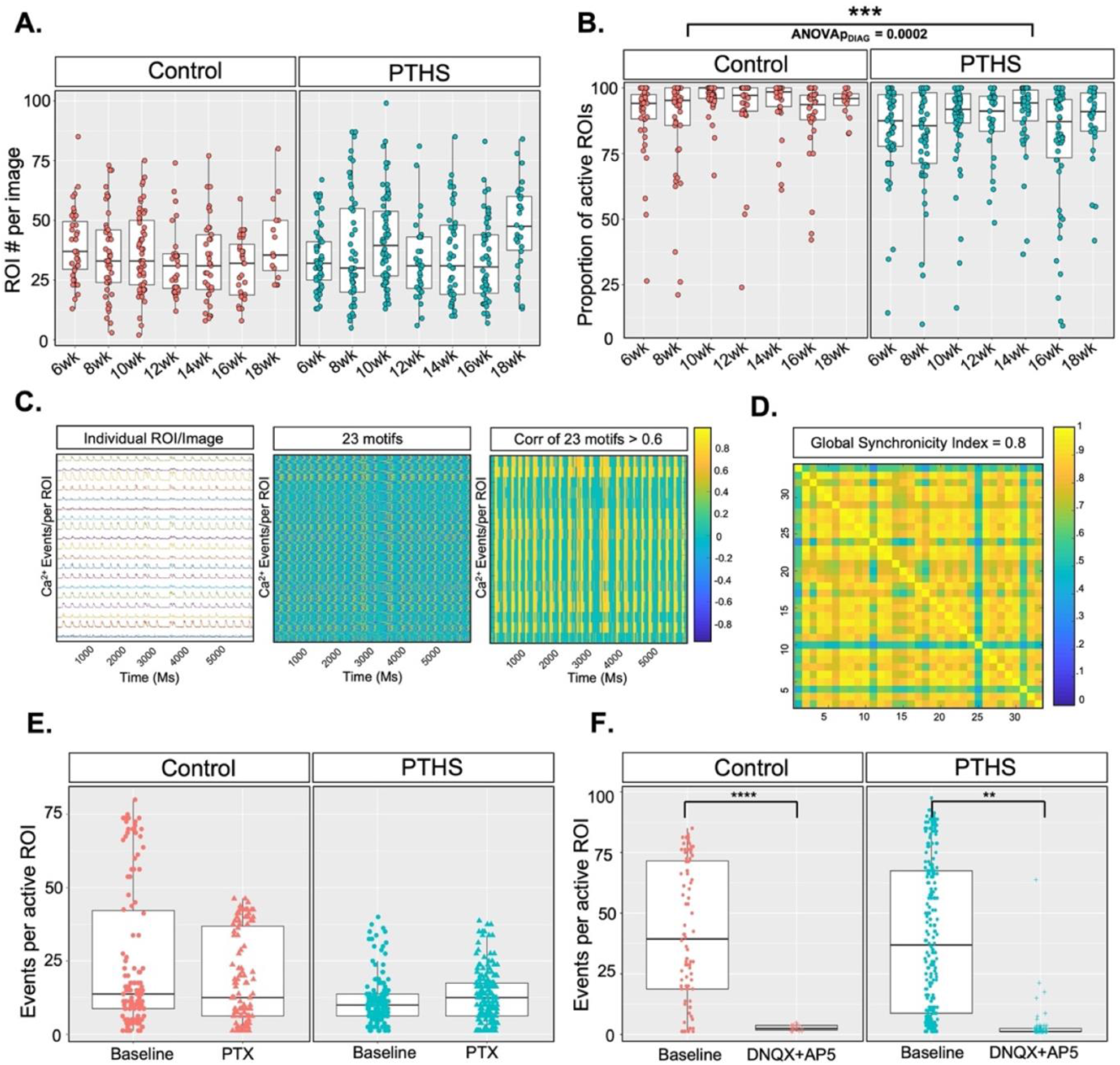
PTHS patient-derived neurons displayed a reduced proportion of active neurons within the network, but equivalent responses to pharmacological blockade of synaptic receptors. **A**. The number of ROIs per image was not different between control and PTHS cultures. **B**. A significant effect of diagnosis was observed for the proportion of active ROIs (diagnosis effect = - 0.1403, p = 0.0002), however the proportion of active ROIs was not different between 6wks and 18wks within controls and PTHS patient lines. Each data point in **A, B**. represents an individual field, two fields were recorded per line, per week, from each differentiation run. **C**. Example Ca^2+^ imaging traces and subsequent plots generated by the CaPTure analysis pipeline to calculate a synchronicity index. Panel 1 shows individual ROI traces representing the change in the normalized fluorescence intensity (ΔF/F, 4 frames/second). Panel 2 shows the motif correlation heatmap with yellow indicating high correlation between a Ca^2+^ event and one of 23 motifs and blue indicating a low correlation between a Ca^2+^ event and the same motif. The turquoise frames represent background activity. Panel 3 shows all frames where the maximum correlation (of 23 motifs) is above a threshold of 0.6 (example from control line N11C1 at 10wks, ROI *N* = 33). **D**. Representative correlation matrix used to calculate the synchronicity index per image which results in a single data point in Figure 3C (example from control line N11C1 at 10wks, ROI *N* = 33). **E**. Group data showing the GABA_A_ receptor antagonist picrotoxin (PTX, 100µM) has no effect on the frequency of network activity in either control or PTHS patient lines in 8wk cultures, a stage when sIPSC frequency was significantly higher in PTHS neurons (Figure 2E). Each data point represents individual ROIs from three control lines and four PTHS lines, with one field recorded, per condition, per line across two differentiations. **F**. Group data showing the co-application of the glutamate receptor antagonists dl-AP5 (100μM) and DNQX (10μM) significantly reduce spontaneous network activity in both control (treatment effect = -31.68, p = 2.03e-08) and PTHS patient lines (treatment effect = -29.059, p = 0.0116), in 18wk cultures when network activity was the highest. Each data point represents individual ROIs from two control lines and two PTHS lines, with one field recorded per condition, per line, across two differentiations.

**Supplemental Figure 9.**
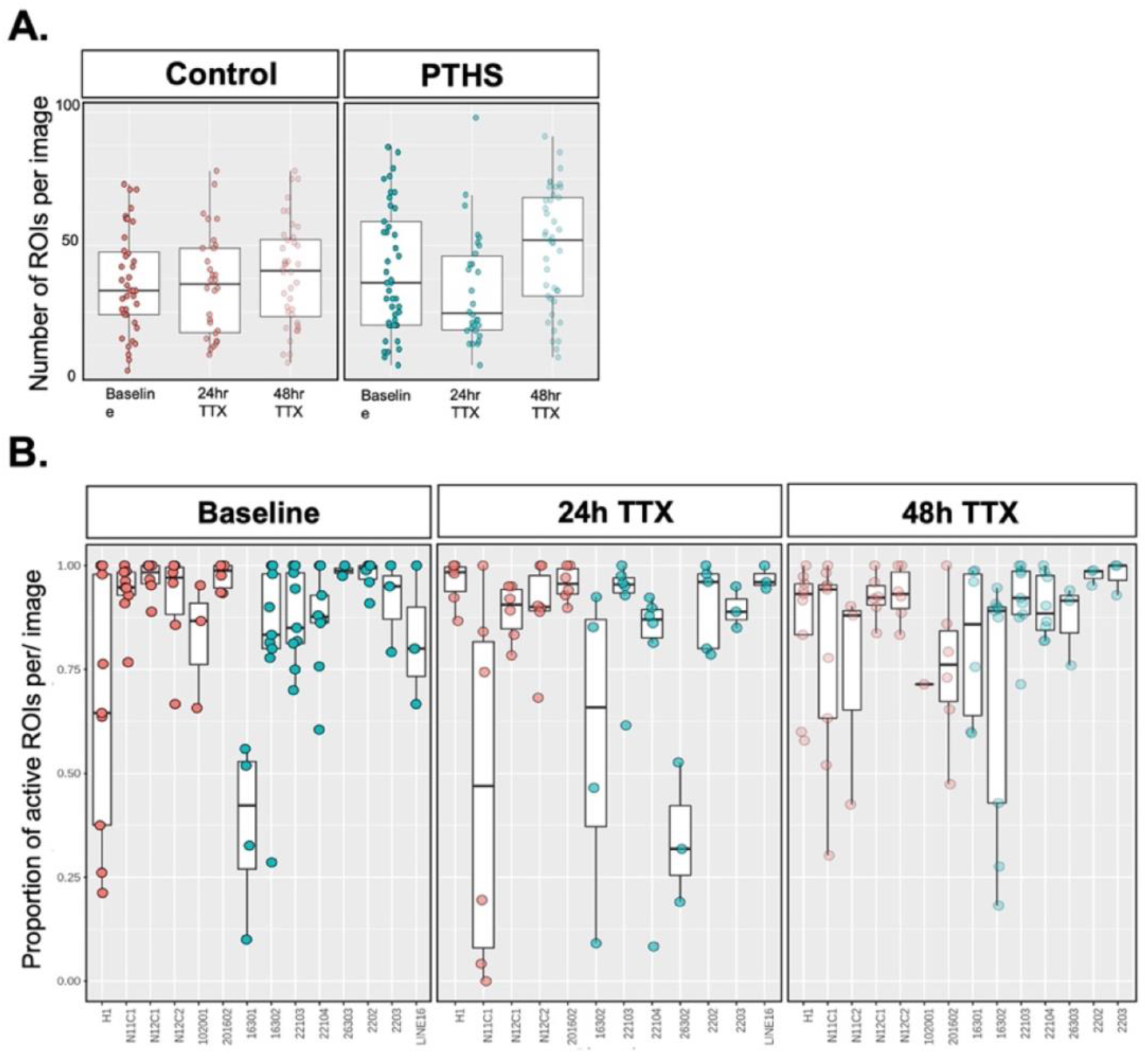
Analysis of neuronal density and within diagnosis variation for homeostatic plasticity experiments. **A**. Summary data showing TTX treatment had no effect on the number of ROIs in an imaging field. **B**. Summary data by line showing TTX treatment had no effect on the proportion of active ROIs per image in either control or PTHS patient neurons. Each data point in A. and B. represents an individual field from the experiment described in Figure 4B.

**Supplemental Figure 10.**
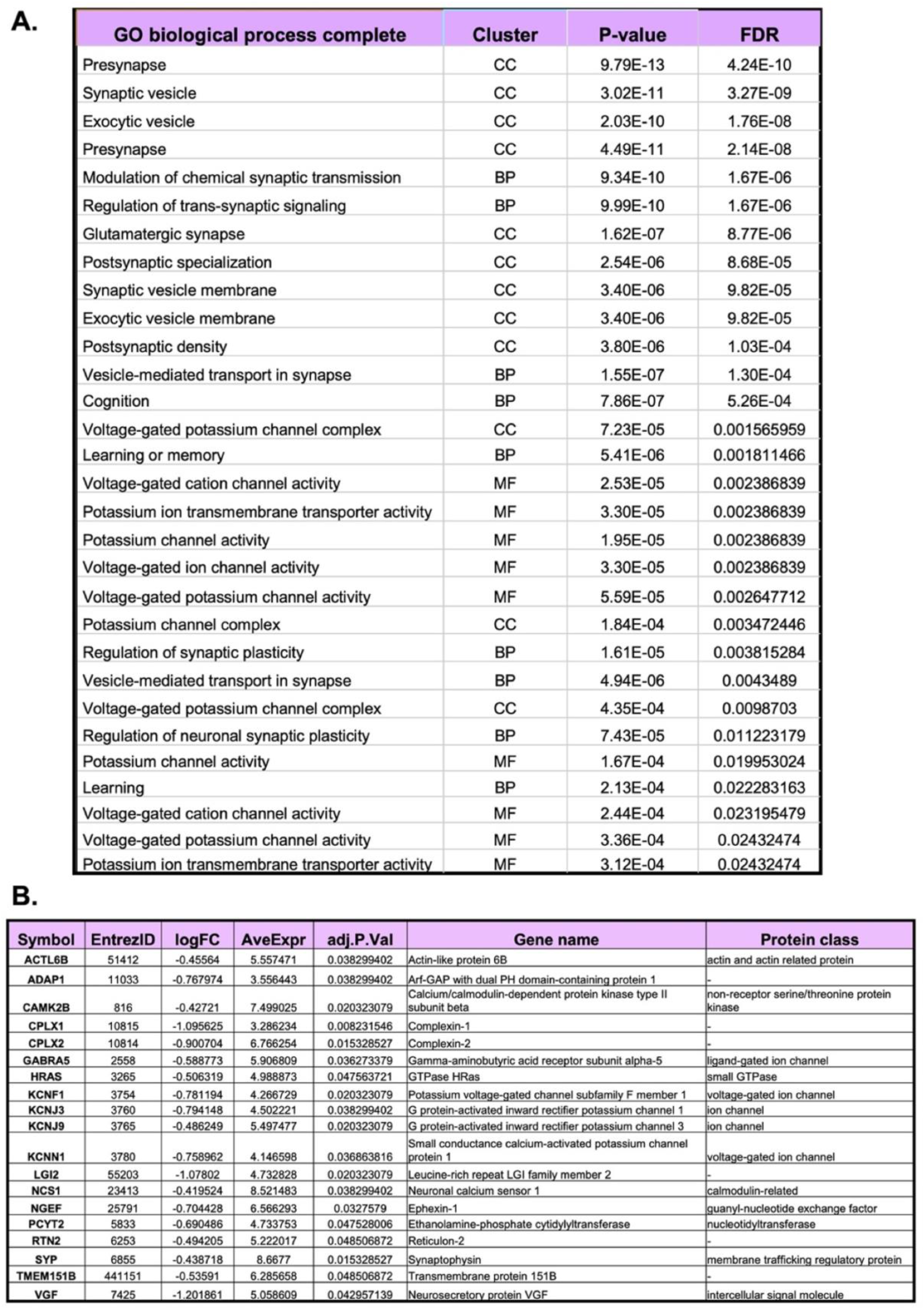
Control neurons demonstrated an appropriate transcriptional response to synaptic upscaling. **A**. Table highlighting the results of gene set enrichment analysis of DEGs (p < 0.005) in control neurons following TTX treatment. **B**. Transcription analysis after homeostatic scaling identified 49 FDR-corrected DEGs in control neurons. Nearly 40% of these genes (19/49) overlapped with a study identifying DEGs after homeostatic upscaling in rat primary cortical cultures. Table of the 19 DEGs found in control neurons that overlap DEGs found in a prior rodent homeostatic scaling study (Benevento et al., 2016).

**Supplemental Figure 11.**
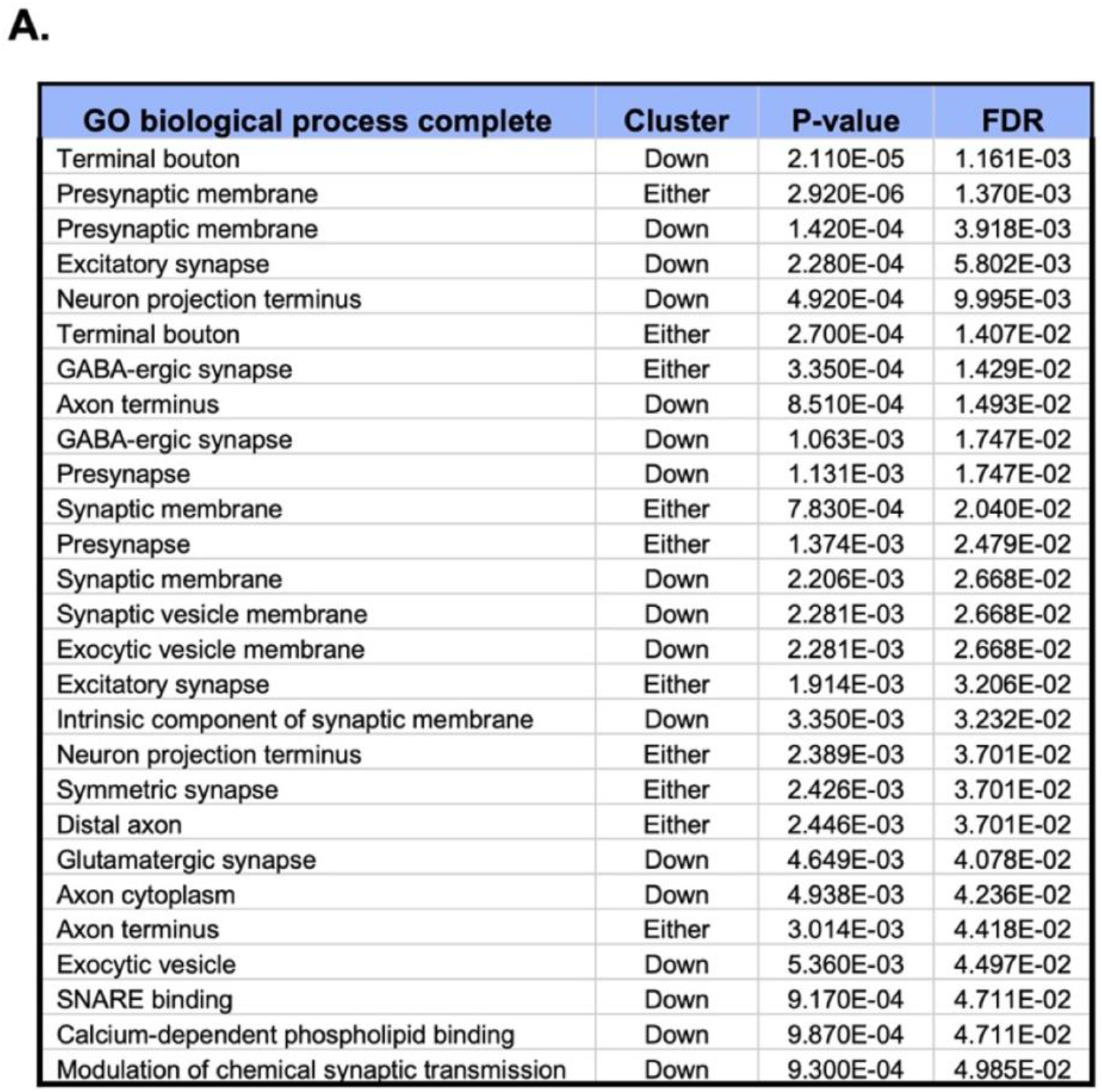
PTHS patient-derived neurons showed transcriptional deficits in genes related to presynaptic function. **A**Table highlighting results of gene set enrichment analysis of differentially expressed genes (p < 0.005) obtained from combined analysis of 8wk plus 18wk PTHS and control neurons.

**Supplemental Figure 12.**
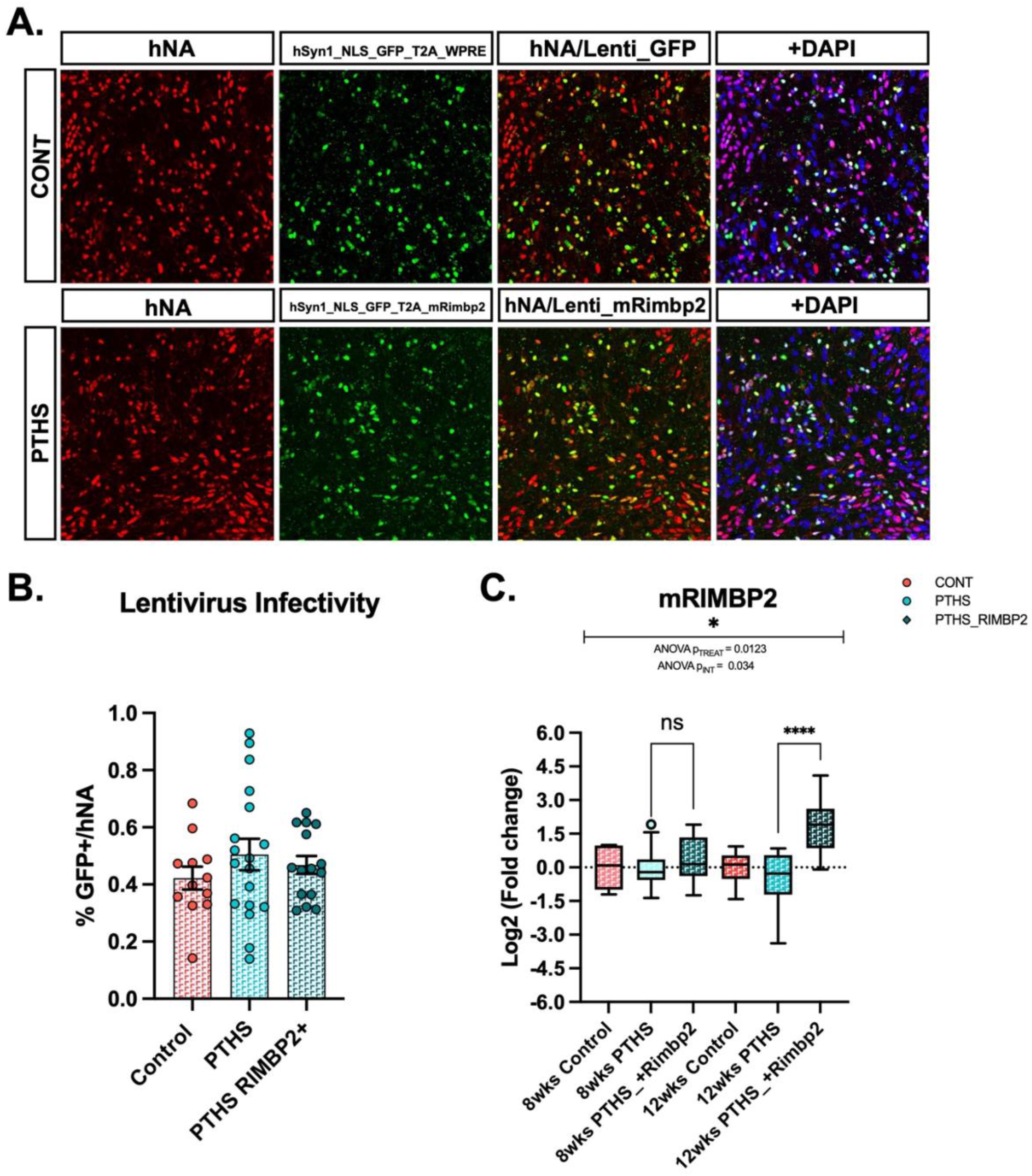
Analysis of lentiviral transduction. **A.** Example ICC images showing colocalization of GFP and hNA after lentivirus transduction in control and PTHS neurons. **B**. Summary plot showing no difference in the infectivity rate of human neurons (%GFP+/hNA); approximately ∼50% of human neurons express hSYN driven GFP+ transduction, independent of construct or diagnosis. Each data point represents the % of GFP+ cells among hNA labeled cells in one field, with at least two fields across a minimum of three experiments for two control lines and four PTHS lines. **C**. Summary plot showing a main effect of treatment on mouse *Rimbp2* expression after viral transduction in PTHS neurons (F (1, 40) = 6.872, p = 0.0123), as well as an age by treatment interaction (F (1,40) = 4.818, p = 0.034). Post hoc analysis reveals the interaction was driven by a significant difference at the later time point, (*t* (1,40) = 5.002, p < 0.0001). Each data point represents mRNA collected from a single well, per condition, with a minimum of one well per line from two independent differentiations, with two control lines and four PTHS lines represented. Values displayed are the log transformation of the delta-delta CT values.

**Supplemental Figure 13.**
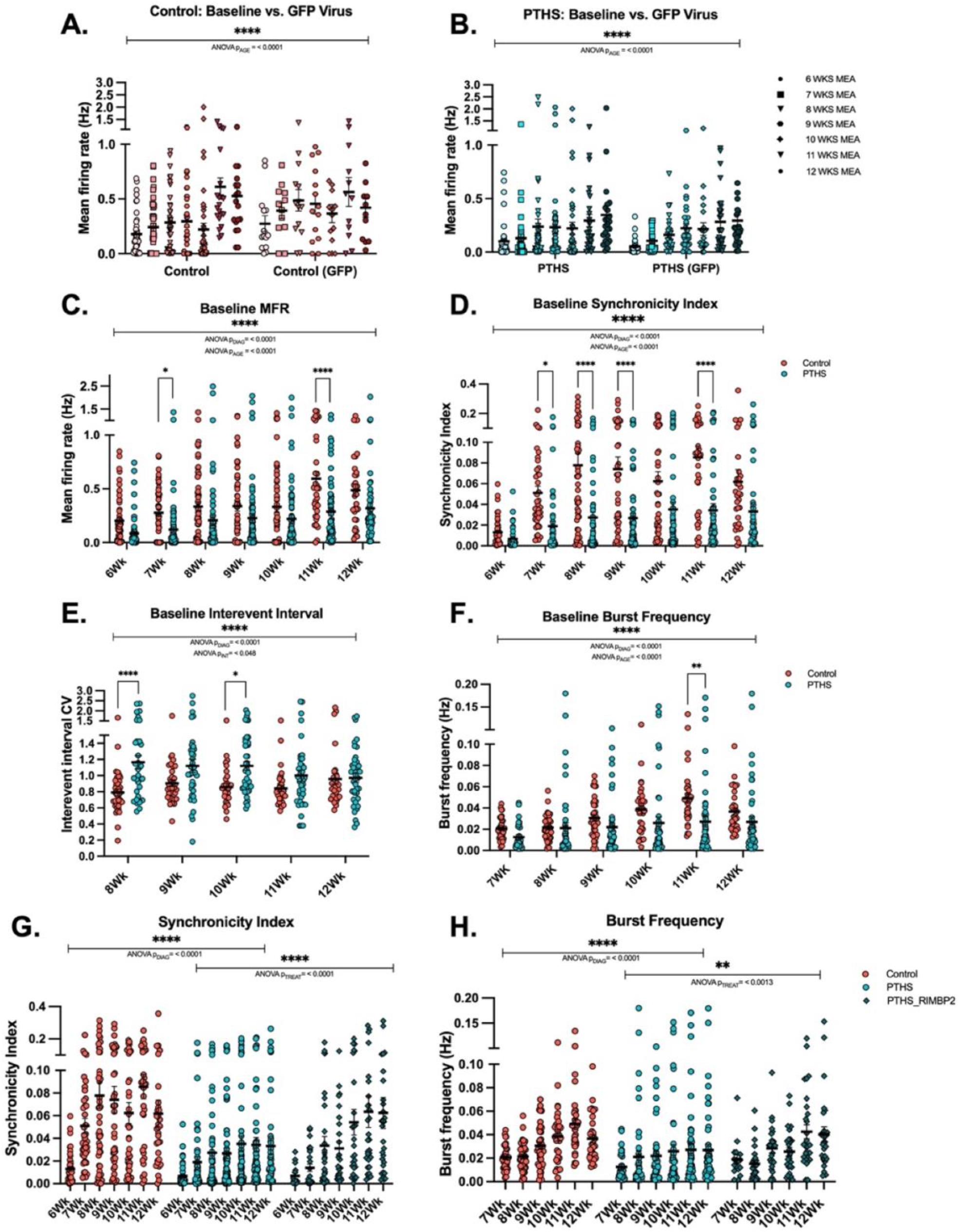
Summary data of MEA recording over development and in response to RIMBP2 expression. **A**. A summary plot showing no effect of GFP viral transduction on the MFR in control neurons between 6 wks and 12wks. **B**. Summary plot showing no effect of GFP viral transduction on the MFR in PTHS neurons between 6wks and 12wks. **C**. Summary plot showing a main effect of diagnosis on the MFR (F (1, 810) = 47.95, p < 0.0001) and a main effect of age (F (6, 810) = 11.21, p < 0.0001) between 6wks and 12wks. **D**. Summary plot showing a main effect of diagnosis on the synchronicity index (F (1, 657) = 77.25, p < 0.0001) and a main effect of age (F (6, 657) = 10.14, p < 0.0001) between 6wks and 12wks. **E**. Summary plot showing a main effect of diagnosis on the interburst interval CV between 6wks and 12wks (F (1, 355) = 28.71, p < 0.0001) and a main interaction effect between diagnosis and age (F (4, 355) = 2.413, p = 0.0488). **F**. Summary data showing a main effect of diagnosis on burst frequency (F (1, 469) = 15.81, p < 0.0001) and main effect of age (F (5, 469) = 6.201, *p* < 0.0001) between 6wks and 12wks. **G**. Summary plot showing a main effect of diagnosis on the baseline synchronicity index (F (1, 565) = 12.98, p < 0.0001) and a main effect of RIMBP2 treatment (F (1, 565) = 8.277, p < 0.0042) between 6wks and 12wks. **H**. Summary plot showing main effect of diagnosis on burst frequency (F (1, 469) = 15.81, p < 0.0001) and a main effect of RIMBP2 treatment (F (1, 409) = 4.060, p = 0.0013) between 6wks and 12wks. Each data point in **A**-**H**. represents a single well, from wells with greater than two active electrodes, from two control lines and five PTHS lines, across three independent differentiations, with two or more wells per condition, per differentiation.

**Supplemental Figure 14.**
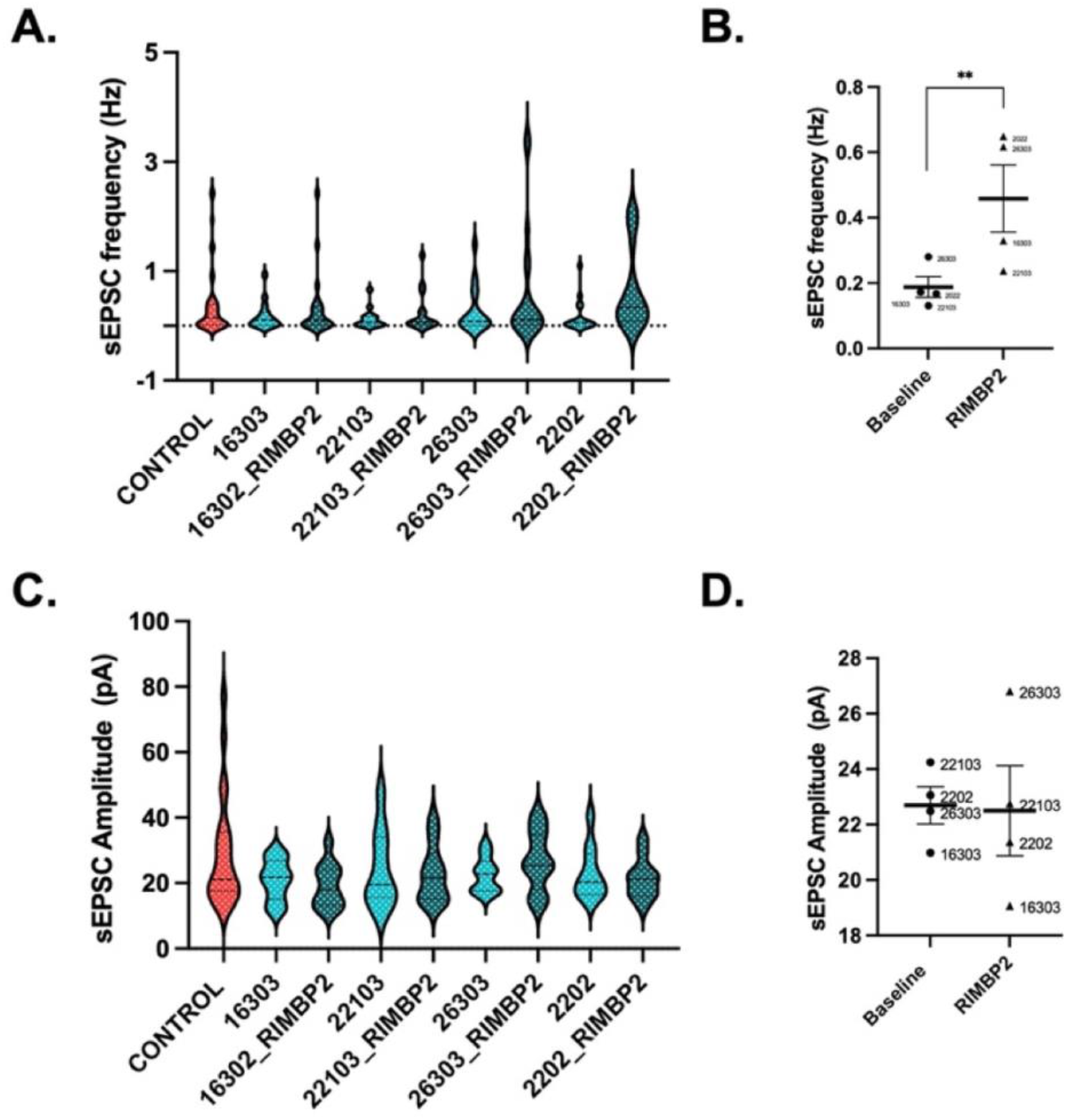
Analysis of within diagnosis variation in RIMBP2 rescue of sEPSC frequency and amplitude. **A**. Quantification of sEPSC frequency by line showed no differences within diagnosis, but a significant RIMBP2 treatment effect (F (1, 129) = 7.176, p = 0.0083). **B**. Summary graph demonstrates sEPSC frequency treatment effects by line. **C**. Quantification of sEPSC amplitude by line analysis showed no differences within diagnosis, or after RIMBP2 treatment. **D**. Summary graph demonstrates no sEPSC amplitude treatment effects by line. Data refers to that described in Figure **5 H**. and **I**.

